# Mechanistic Insights into the inhibition of *Plasmodium falciparum* DNA gyrase A by withanolides derivatives through integrated computational analysis

**DOI:** 10.1101/2025.09.30.679425

**Authors:** Bini Chhetri Soren, Soujanya J. Vastrad, Syed Junied, Diya Srikanth, Aparna Chimalamari, BJ Mahender Kumar, Jagadish Babu Dasari

**Affiliations:** Department of Research and Development, Quanton BioLife Sciences, Ghatsila, Jharkhand, India; Department of Pharmacy Practice, Faculty of Pharmacy, M.S. Ramaiah University of Applied Sciences, New BEL Road, M.S.R. Nagar, Bengaluru, India; University of Milan, Department of biosciences, Via Festa del Perdono 7, Milano, Italy; Department of Pharmacy practice, Faculty of Clinical Pharmacy, SAC college of Pharmacy, BG Nagara, Karnataka, India

**Keywords:** Malaria, *Plasmodium falciparum* DNA Gyrase A, Withanolide, Molecular docking, Binding energy, Molecular dynamic simulations

## Abstract

Malaria is a fatal disease affecting millions of people worldwide, primarily due to infection by *Plasmodium falciparum*. The emergence of multidrug-resistant parasite strains has necessitated the exploration of novel therapeutic targets, among which DNA gyrase represents a unique and underexploited enzyme in the parasite’s replication machinery. *Plasmodium falciparum* DNA gyrase A (pfDNA gyrase), an essential topoisomerase II that is not present in humans, has been identified as a promising target for antimalarial drug development. Present study deals with a structure based computational approach to characterize the binding mechanism and dynamic stability of three bioactive withanolide derivatives (D, E, and O) against pfDNA gyrase. Molecular docking revealed high binding affinities for withanolide D (-9.14kcal/mol), E (-9.73kcal/mol), and O (-9.00kcal/mol), with interactions mediated through key catalytic residues such as GLU648, LYS647, and TRY590 via hydrogen bonding and hydrophobic contacts. Stability of the ligand-protein complexes was further assessed through molecular dynamics simulations, where analyses of RMSD, RMSF, radius of gyration (Rg), and solvent accessible surface area (SASA) analysis confirmed the structural integrity and compactness of the complexes, notably withanolide O exhibited the most favorable dynamic profile, whereas withanolide E induced confirmational rigidity. MM/GBSA calculations are further supported by showing the lowest binding free energy for withanolide O and E (ΔG_bind_ =-20.89 and -20.22 kcal/mol). The ADME studies showed favorable pharmacokinetic and physiochemical properties of three ligands. Collectively, these findings highlight the potential of withanolide derivatives as promising inhibitors of pfDNA gyrase, thereby paving a way for the foundation of future antimalarial drug development.

## 1. Introduction

Malaria, a parasitic disease caused by infections related to plasmodium, remains a global medical concern and challenge. It is caused by five species of protozoan parasite, namely *Plasmodium falciparum, Plasmodium vivax, Plasmodium ovale, Plasmodium malariae, and Plasmodium knowlesi* [1,2]. Amongst all these malaria causing agents, *Plasmodium falciparum* infection remains the most lethal, resulting in the most of the death cases, whereas the others cause a slightly lesser effect. The parasite’s complex life cycle that involves a human host and mosquito vector makes it even more difficult for the eradication effort. In addition, the rise of drug-resistant strains has resulted in treatment becoming even more challenging. Thus, increasing the urgency to develop new therapeutic targets. One such promising target is the DNA gyrase enzyme present in the apicoplast of *Plasmodium falciparum* [3]. Furthermore, the apicoplast can serve as a significant target, as this is a unique organelle present in protozoan parasites but absent in human cells, thereby rendering an opportunity for selective drug development and also minimizing off target effects [4].

DNA gyrase is a topoisomerase type II responsible for the DNA -binding, cleavage, and religation of negative supercoils into DNA, thereby playing a vital role in biological processes. These enzymes are categorized into two groups, Type I and Type II, based on their DNA strand breaks. In prokaryotes, Type IIA is responsible for catalyzing ATP-dependent DNA supercoiling by A_2_B_2_ heterotetramer complex, which binds to positively supercoiled DNA and uses ATP hydrolysis to cause negative supercoiling. Moreover, the N-terminal domain of DNA gyrase A is responsible for breaking and religating the DNA, whereas the C-terminal domain is associated with wrapping the DNA. Additionally, N-terminal domain in gyrase B is linked to ATPase activity, while the C-terminal domain shows DNA binding and interaction with gyrase A [5, 6]. Thereby demonstrating that both subunits of DNA gyrase are vital for DNA topology. Although pfDNA gyrase A has been the subject of extensive research, the association of its biological function and its reactivity towards *Plasmodium falciparum* DNA gyrase A (pfDNA Gyrase A) inhibitors for blood-stage malaria parasite remain ambiguous [6]. Furthermore, the mechanistic insights against pfDNA gyrase A domain (PDB:6CA8), which is responsible for cleavage and relegation remains unclear. Additionally, the disruption of pfDNA gyrase A using CRISPR/CAS9 has revealed its sensitivity towards DNA topology within the apicoplast genome [7].

Currently, various inhibitors of DNA gyrase have been identified and synthesized; these are classified according to their chemical structure. However, due to the significant rise in antibiotic-resistant pathogens, there is an urgent need to explore novel natural derivatives as potential inhibitors [8]. Therefore, it is critically important to investigate new potential drugs that can specifically target DNA gyrase, particularly pfDNA gyrase, to address the current challenges. One such natural derivative could be bioactive compounds extracted from the plant *Withania somnifera* (W.somnifera), which has been previously shown to be effective in antibacterial [9, 10], anti-cancer, anti-immunomodulator [11, 12, 13], anti-inflammatory [14, 15, 16], and anti-parasite properties [17,18]. Withanolide derivatives, which belong to the Solanaceae family, consist of C28 steroids featuring an ergostane skeleton. In this structure, C22 or C23 and C26 are oxidized to form a delta or gamma lactone ring; further addition of rings to the fundamental skeletal structure with various chemical groups contributes to the development of numerous molecules that are classified as withanolides [19, 20]. The present study aims to investigate the inhibitory properties of twelve withanolides, out of these three withanolides, namely withanolides D, E, and O, have been extensively studied for their mechanism of action against pfDNA gyrase A (PDB:6CA8) and their structural interactions.

## 2. Methodology and Materials

### 2.1. Protein Structure Preparation

The crystal structure of *Plasmodium falciparum* DNA gyrase (pfDNA gyrase) was retrieved from the Protein Data Bank (PDB ID: 6CA8) and contained unsolved regions due to missing amino acids residues loop segments. To generate a complete and structurally accurate model, Modeller v10.6 [21] was employed to reconstruct the missing residues through comparative homology modeling. The sequence alignment was performed against the full-length reference sequence, and ten models were generated. The model with the lowest DOPE (Discrete Optimized Protein Energy) score was selected for further analysis and visualization, as illustrated in supplementary Fig. 1(a). Prior to docking, the protein was processed using Auto Dock Tools v4.2.6 [22], where all non-standard residues, water molecules, and pre-bound ligands were removed. Polar hydrogens were added, Gasteiger-Marsili charges were assigned, and the file was saved in PDBQT format for molecular docking.

### 2.2. Ligand Preparation

The *W.somnifera* derivatives of twelve withanolides ligand were selected for molecular docking studies. These compounds were retrieved from Indian Medicinal Plants Databases (IMPD) [23] and PubChem in PDB format. Energy minimization and geometry were carried out by using Avogadro version 2.0.6. Each ligand was added, non-polar hydrogen bonds were merged, Gasteiger-Marsili charges were added, atoms were adjusted according to AutoDock atom types, and rotatable bonds were assigned. Then, ligands were converted into PDBQT format using AutoDock Tools v1.5.7.

### 2.3. Molecular Docking Studies

Molecular docking simulations were performed using AutoDock Tools v1.5.7. The 2.33Å of X-ray crystallographic structure of pfDNA gyrase (6CA8) was selected as the binding pocket for molecular dynamics screening. Stereochemical quality and Ramachandra plot was evaluated using PROCHECK [24], exhibited with 635(92.6%) residues are favored regions and a further 50 residues (7.3%) allowed regions, and 1 (0.1%) at the outlier shown in supplementary Table 1. The docking grid box was centered around the active site of pfDNA gyrase, determined by using AutoDock Tools v1.5.7. For each ligand, the best pose was selected based on the lowest binding energy score and visual interaction analysis. Detailed results for all twelve ligands were summarized in supplementary Table 2.

### 2.4. Visualization and Interaction Mapping

The docked complexes were visualized and analysed using Discovery Studio Visualizer v2024 (BIOVIA, Dassault Systems) [25]. For each ligand–protein complex, ribbon diagrams, the two-dimensional (2D) interaction maps, and three-dimensional (3D) stick models were generated. Hydrophobic surface maps were colored from hydrophilic in blue and hydrophobic in brown. Hydrogen bond donor and acceptor surfaces were mapped in magenta and green, respectively. Interaction types were identified, including conventional hydrogen bonds, π-alkyl, alkyl, and unfavorable electrostatic interactions.

### 2.5. Molecular dynamics Simulations and trajectory analysis

The microscopic stability of proteins and their complexes with ligands is evaluated through dynamics, a sophisticated automated simulation technique. This evaluation encompasses aspects such as structure, function, fluctuations, interactions, and overall behaviour. The dynamics of the three complexes were analysed utilizing GROMACS version 2025.1 for molecular dynamics calculations [26] under Linux platform. The CHARMM General Force Field was employed to parameterize the protein components, while ligand topologies were generated using the Swiss Param server [27]. The structures underwent vacuum minimization 2500 times via steepest descent to mitigate steric conflicts. Subsequently, Simple Point Charge (SPC) water model was used to solvate the structures. To achieve electrical neutrality, Na^+^ and Cl^-^ ions were introduced, facilitated by the gmx genion tool. Following minimization, the molecular dynamics simulations progressed into production phases, specifically NVT and NPT. These two phases were crucial for system balancing. Initially, a 100 picosecond NVT equilibration was conducted to stabilize the number of particles, volume, and temperature, with the objective of elevating the system temperature to 300 K. The second phase involved a meticulous 100 picosecond NPT equilibration to ensure uniformity in temperature, pressure, and particle count, which was vital for maintaining system density and pressure. During the simulations, the positioning of the protein group was constrained by bond limitations on all connections. The entropy of the system decreased as NVT and NPT conditions restricted the movement of water molecules surrounding the protein, allowing them to relax. The Parrinello-Rahman barostat [28] method and v-rescale thermostat were calibrated for a duration of 100 picoseconds. To enforce covalent bond constraints, the Linear Constraint solver application was utilized. The advanced Particle-Mesh Ewald (PME) method managed chemical bond interactions enabling 100 ns of production for each system. The MD analysis were analyzed by plotting Root Mean Square Deviation (RMSD), Root Mean Square Fluctuation (RMSF), and Radius of gyration (Rg) through qtgrace tool.

### 2.6. MM/GBSA Binding Free Energy

To evaluate the binding free energy of the ligand to protein, we conducted Molecular Mechanics/Generalized Born Surface Area (MM/GBSA) calculations utilizing the gmx_MMPBSA tool version 1.6.3, which integrates GROMACS with AmberTools 2022 for comprehensive end-state free energy analysis [29]. A total of 1000 frames extracted from the last 50 ns of the production run were analysed. The electrostatic energy (EEL), polar solvation energy (EGB), and nonpolar solvation energy (ESUR). The total binding free energy (ΔG_bind_) was computed by summing these contributions, providing insight into the relative stability and affinity of each protein-ligand complex. This methodology dissects the binding energy (ΔG_bind_) into components of molecular mechanical energy (van der Waals and electrostatic interactions) and solvation energies (polar and nonpolar), shown in the following equation:

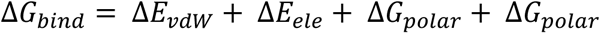

In this investigation, the Generalized Born (GB) implicit solvent model was employed to assess polar solvation energy, specifically using parameters aligned with the GB-Neck2 model (igb=8), while the non-polar contribution was derived from solvent accessible surface area through Linear Combinations of Pair Wise Overlaps (LCPO) method. Entropy effects were disregarded, a common practice in comparative binding affinity evaluations [30]. A total of 2501 snapshots were collected at uniform intervals from the 10 ns production trajectory (MD_center.xtc) of the protein-ligand complex. The calculations were performed on three distinct systems: the complex, the receptor (protein alone), and the ligand. The final ΔG_bind_ was determined by using the formula:

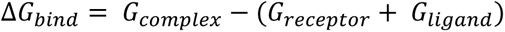

### 2.7. Principal Component Analysis (PCA)

PCA was performed to evaluate the primary movements within the protein-ligand complex throughout the molecular dynamic simulation. This technique effectively reduces the dimensionality of atomic fluctuations by diagonalizing the covariance matrix derived from positional deviation after least squares fitting to a reference structure, resulting in the extraction of eigenvectors (principal components) and eigenvalues (indicating their amplitudes) [31, 32]. The GROMACS 2025.1 software was employed for trajectory pre-processing and the execution of PCA. The atomic positional co-variance matrix was constructed using the *gmx covar* command applied to the Cα atoms of the pfDNA gyrase, which yielded the corresponding eigenvectors and eigenvalues. Following this, the *gmx aneig* tool was utilized to examine the first three principal components in relation to RMS fluctuation, contributions of eigenvectors, and 2D trajectory projections. All PCA visualizations were created using Xmgrace2025.1. The eigenvectors illustrate the directions of motion within the conformational space, while the eigenvalues reflect their relative amplitudes [33].

### 2.8. Dynamic Cross-Correlation Matrix (DCCM)

To investigate the correlated motions at the residue level within protein-ligand complexes, a DCCM analysis was conducted utilizing the Cα atoms of the protein derived from the 200 ns molecular dynamics (MD) trajectories. The MD trajectories underwent processing through MD analyses in Python 3.9, where displacement vectors from mean positions were calculated for each residue overtime. The normalized correlation coefficients between pairs of residues were determined using the formula:

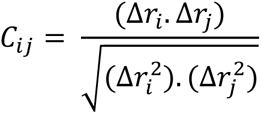

The values of *C_ij_* range from +1, indicating fully correlated motion, to -1, representing entirely anti-correlated motion, with a value of 0 signifying no correlation. The resulting matrices were visualized through seaborn, offering valuable insights into the allosteric communication and long-range interactions influenced by ligand binding, as established in prior research [34, 35]

### 2.9. Free Energy Landscape (FEL)

FEL analysis was performed to assess the conformational energetics and stability of the three complexes. The trajectory files generated from 250 ns molecular dynamics simulations served as the input for PCA, from which the first two eigenvectors (PC1 and PC2) were chosen as reaction coordinates [36]. The free energy associated with each conformation was computed using the Boltzmann relation:

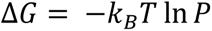

where ΔG is the free energy, kB denotes the Boltzmann constant (1.380649 × 10⁻²³ J/K), T indicates the temperature (298.15 K), and lnP is the normalized probability distribution of the conformations in the PC1–PC2 space [37]. The 3D FEL was illustrated using matplotlib.pyplot.plot [38] surface to depict local energy minima and transition states across a complex conformational landscape. This methodology facilitates the identification of predominant low-energy basins and metastable states, a technique commonly utilized in studies of protein dynamics [39, 40].

### 2.10. Analysis of ADME properties

Subsequent to the docking analysis, the withanolide ligands underwent the SwissADME web tool [41], which is freely available at http://www.swissadme.ch, and the pkCSM server https://biosig.lab.uq.edu.au/pkcsm was also used for detailed analysis of ADME properties of ligands [42]. These applications assess vital pharmacological properties, including lipophilicity, water solubility, drug likeness, bioavailability, and numerous other chemical features that are crucial for drug discovery. The SMILES structures of the ligands were uploaded to the server to retrieve all pertinent ADME attributes.

## 3. Results

### 3.1. Molecular docking and interaction analysis

To elucidate the binding modes and interaction mechanisms of withanolide derivatives, molecular docking was performed using Autodock Tools version 1.5.7, targeting the DNA gyrase of *Plasmodium falciparum* (PDB: 6CA8). The docked complexes of withanolide derivatives, as shown in supplementary Table 2. Among these, withanolide D, E, and O were evaluated for hydrogen bonding, hydrophobic contacts, and structural complementarity within the enzyme’s binding pocket. Withanolide D, with a binding affinity of -9.14 kcal/mol, docked into the catalytic cleft of pfDNA gyrase and adopted a stable orientation supported by key interactions shown in Fig.1a-e. Notably, it formed conventional hydrogen bonds with GLY409, GLU411, GLY353, and LYS280, as well as π-alkyl interactions with TRP488, LYS486, and PHE332. An unfavorable acceptor-acceptor contact was noted with GLY353, indicating a close spatial proximity of electron-rich regions. The hydropathy surface map in Figure 1d revealed a moderately polar environment, with pockets of hydrophobicity aligning with the ligand’s nonpolar moieties. Additionally, the hydrogen bonding surface confirmed complementary donor and acceptor regions along the binding interface, indicating a stable yet moderately flexible binding profile. Withanolide E, exhibiting the highest binding affinity among the tested derivative at -9.73 kcal/mol displayed a tightly embedded pose within the pfDNA gyrase active site, forming a dense bonding network involving GLU648, GLU649, and LYS647, as shown in Figure 2a-e. Polar contacts were established with TYR590, LYS384, and LYS597, enhancing electrostatic complementarity. These interactions contributed to a highly stabilized docking confirmation. The hydrophobicity map shows a predominantly polar environment, while π-alkyl and alkyl contacts with TYR590 and LYS384 show a hydrophobic balance seen in Fig. 2a. The hydrogen bond mapping supported a multivalent interaction model, as depicted in Figure 2e. Whereas in withanolide O, with a binding affinity of -9.00 kcal/mol, occupied a semi-enclosed grove within pfDNA gyrase and was anchored through both conventional and carbon-hydrogen bonds, interacting with the residues included LYS647, ASN586, and LEU383 as shown in Fig. 3a-e. The binding pose suggested a tight fit, stabilized by dual-mode polar interactions and alkyl contacts. The hydrophobic mapping and hydrogen bond surface revealed the polar dominance in the binding cavity, complemented by neutral hydrophobic zones engaging with the ligand’s steroidal core and accurate spatial alignment of donor and acceptor atoms, as illustrated in Fig. 3d and 3e.

**Figure 1:**
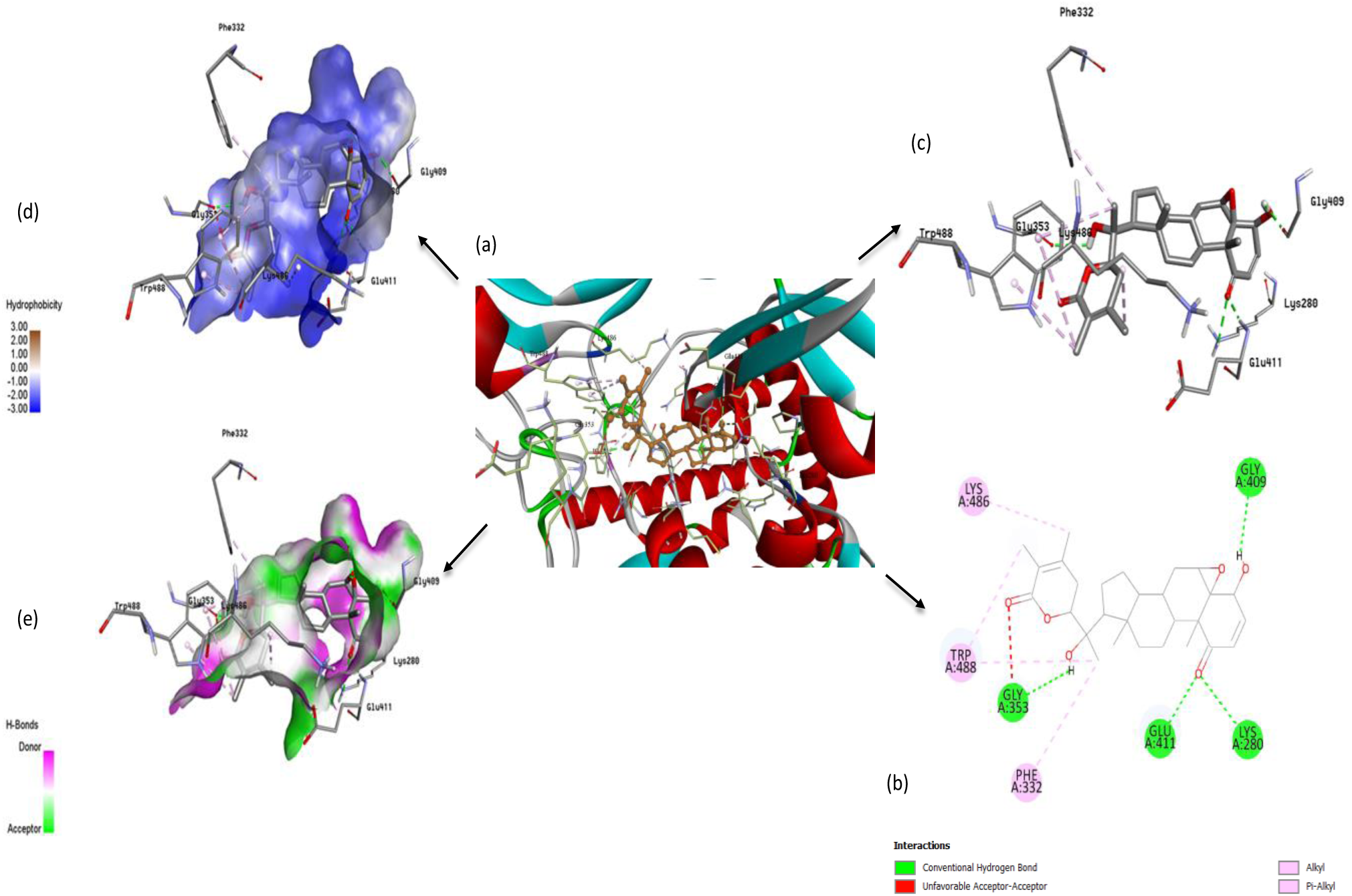
Binding interactions of withanolide D against pfDNA gyrase (PDB:6CA8). Panel (a) shows the ribbon representation of the protein with helices in red, β-sheets in cyan, and loops in grey, highlighting the ligand in orange sticks positioned within the active sites. Panel (b) details the two-dimensional (2D) interactions diagram of ligand-residue contacts, where green lines represent conventional hydrogen bonds, pink lines indicate π-alkyl interactions, and red lines denote unfavorable acceptor-acceptor contacts, illustrating the complexity of the binding environment. Panel (c) provides stick models of the ligand interacting amino acids with dashed lines depicting hydrogen and hydrophobic interactions. Panel (d) shows hydrophobicity surface maps, the binding pocket from hydrophilic (blue) to hydrophobic (brown) regions. Panel (e) illustrates the spatial distribution of hydrogen bond donor (magenta) and acceptor (green) regions surrounding the ligand-binding site.

**Figure 2:**
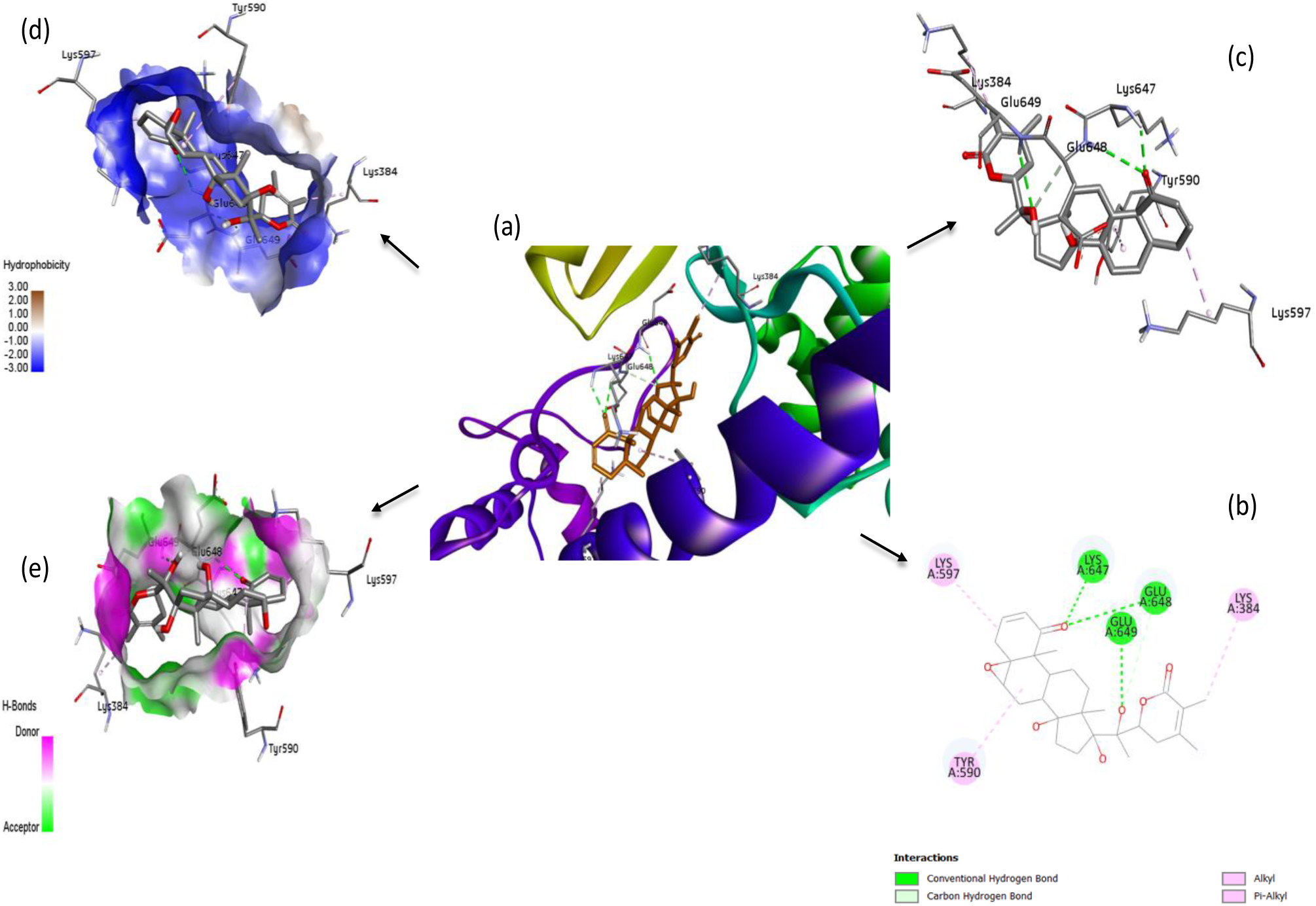
Molecular docking visualization of withanolide E bound to Plasmodium falciparum DNA gyrase (PDB: 6CA8). Panel (a) depicts ribbon representation of the pfDNA gyrase highlighting withanolide E in orange stick model within active site. The α-helices in purple, β-sheet in green/cyan and loops in yellow, illustrates the spatial positioning of the ligand. Panel(b) the 2D interaction diagram shows the ligand-residue contacts, where green lines indicate conventional hydrogen bonds and pink lines represent π-alkyl and alkyl interactions. Panel (c) presents a stick model of the ligand and interacting amino acids, with dashed lines illustrating the non-covalent interactions, hydrogen bonding and hydrophobic contacts within the active site. Panel (d) hydrophobicity molecular surface of the binding pocket ranging from hydrophilic (blue) and hydrophobic (brown). Panel (e) provides a surface representation of hydrogen bond donor (magenta) and acceptor (green) in regions surrounding the ligand.

**Figure 3:**
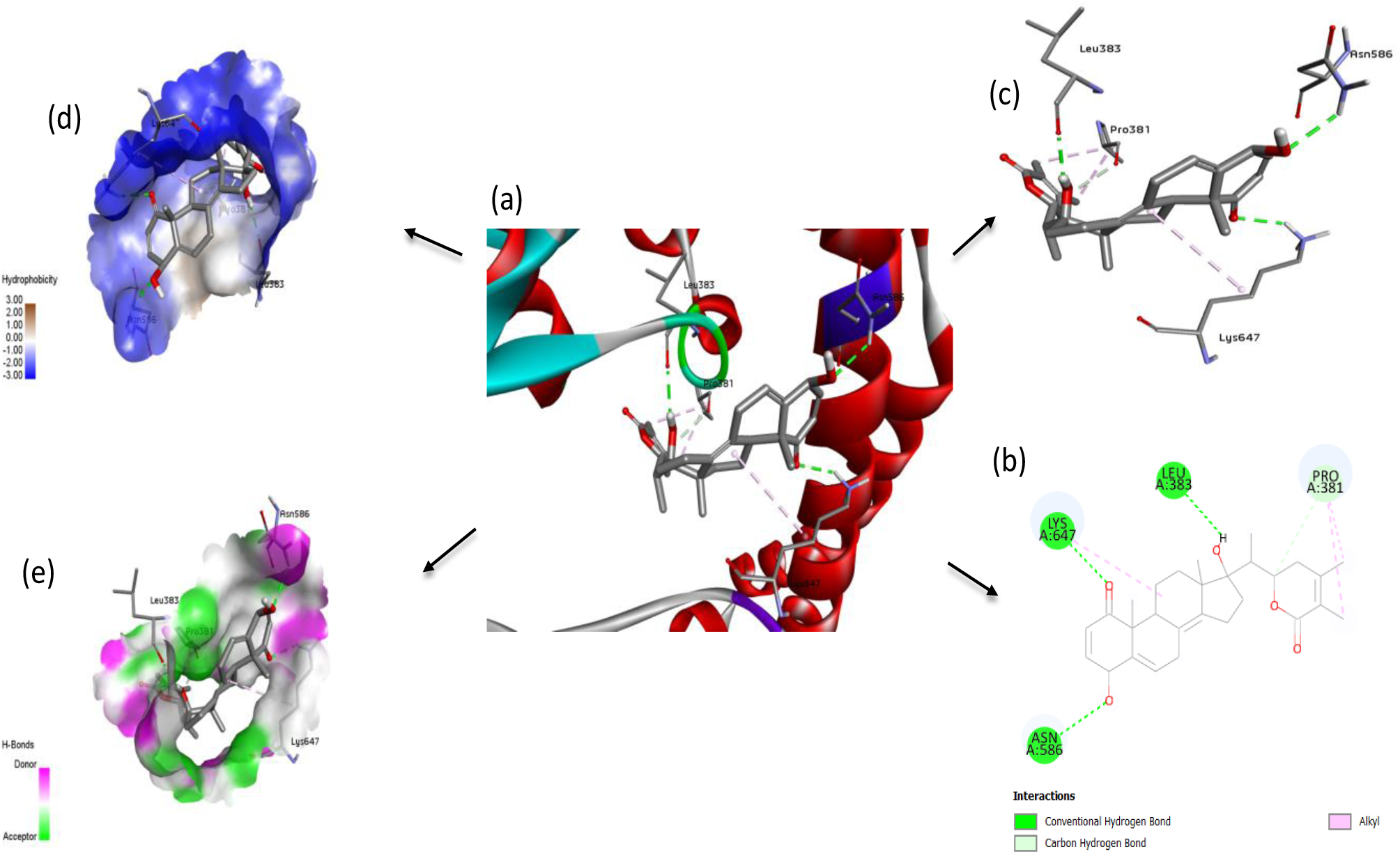
Binding interactions of pfDNA gyrase (PDB:6CA8) with withanolide O. Panel (a) presents the ribbon representation of the pfDNA gyrase protein, with α-helices in red colored and β-sheets in cyan, and withanolide O in grey stick model position within the active sites. Panel (b) show 2D interactions of the ligand-residue contacts. Green lines represent conventional hydrogen bonds, while pink lines represent alkyl and carbon hydrogen bond interactions. Panel (c) provides a close tick model of withanolide O and the surrounding amino acid residues. Green and grey dashed lines illustrate the hydrogen and hydrophobic contacts. Panel (d) displays the molecular surface of the binding colored by hydrophobicity, with blue indicating hydrophilic regions and brown indicating hydrophobic areas. Panel (e) illustrates the hydrogen bond donor (magenta) and acceptor (green) regions on the protein surface surrounding withanolide O.

### 3.2. Root Mean Square Deviation (RMSD) analysis

The RMSD of the Cα atoms of Apo pfDNA gyrase in the protein exhibited a relatively stable pattern throughout the simulation, oscillating between approximately 0.06 and 0.15nm, with an average value nearing 0.10 nm. In contrast, the ligand withanolide D demonstrated negligible deviation, with its RMSD predominantly ranging from 0.03 to 0.08 nm, indicating a strong binding stability and limited conformational changes. The ligand’s consistent trajectory throughout the simulation highlights a robust and stable interaction with the protein, as illustrated in Figure 4(a). The withanolide E complex, on the other hand, showed increased fluctuations in the Cα pfDNA gyrase atoms, varying between 0.25 and 0.65nm, suggesting a higher degree of global flexibility within the protein structure. Throughout the majority of the trajectory, the ligand’s RMSD remained remarkably stable, fluctuating around 0.04 to 0.06 nm. However, a notable spike was observed between 150 and 180 ns, where the RMSD reached approximately 0.18 nm. This spike likely indicates a temporary conformational change or rearrangement of interactions, with the subsequent return to the initial RMSD values suggesting that the ligand successfully reverted to its original conformations as depicted in Figure 4(b). Lastly, the withanolide O complex demonstrated the most extensive range of RMSD, spanning from 0.35 to 0.72 nm, with a noticeable increase in fluctuations observed towards the end of the trajectory. Conversely, the ligand withanolide O exhibited a consistently low RMSD, ranging from 0.05 to 0.15 nm, reinforcing the notion of stable binding within a relatively flexible protein environment. This observation may indicate that the ligand remains stable in the binding site even as the protein navigates broader conformational space, as shown in Figure 4 (c).

**Figure 4:**
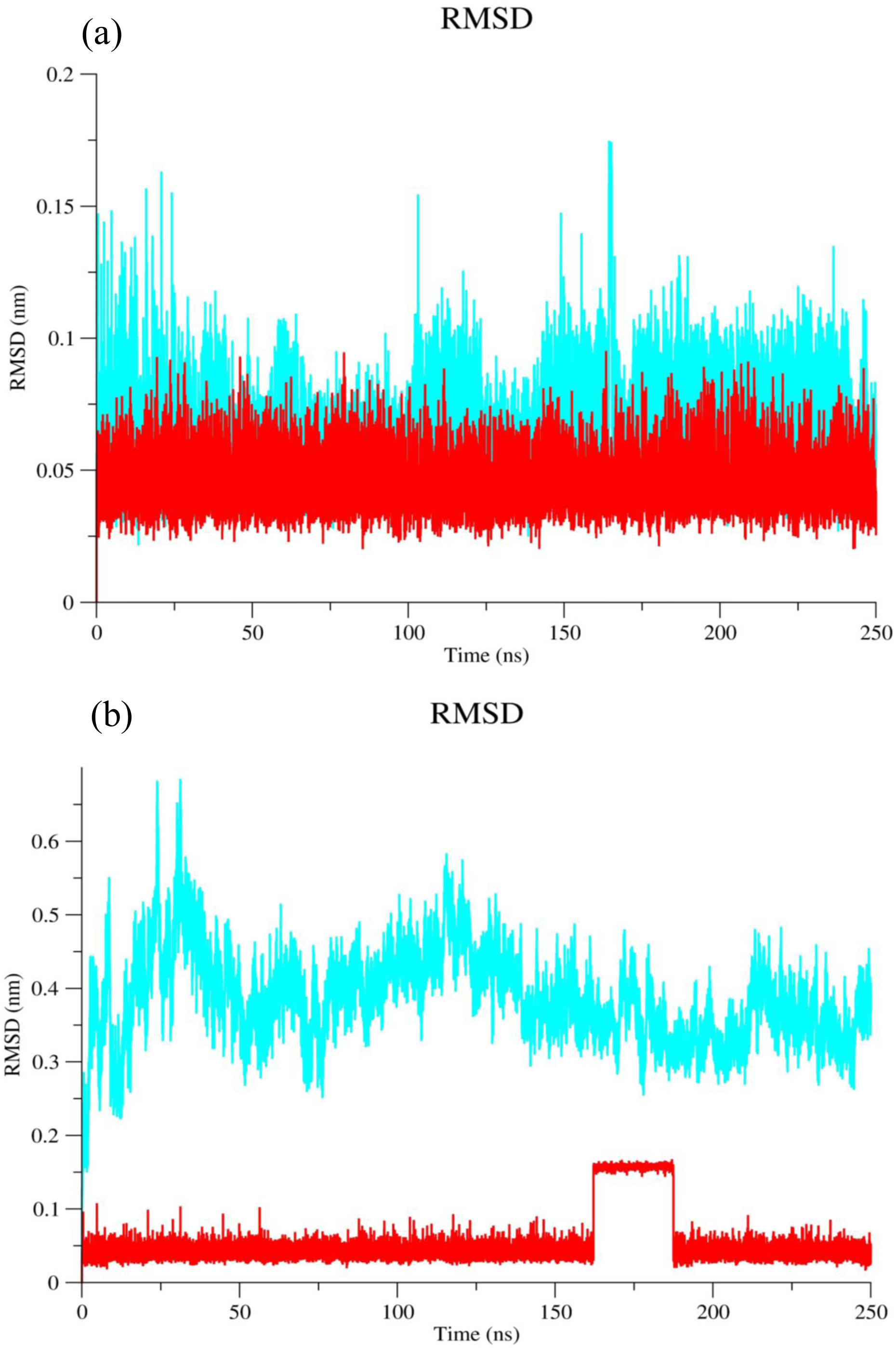
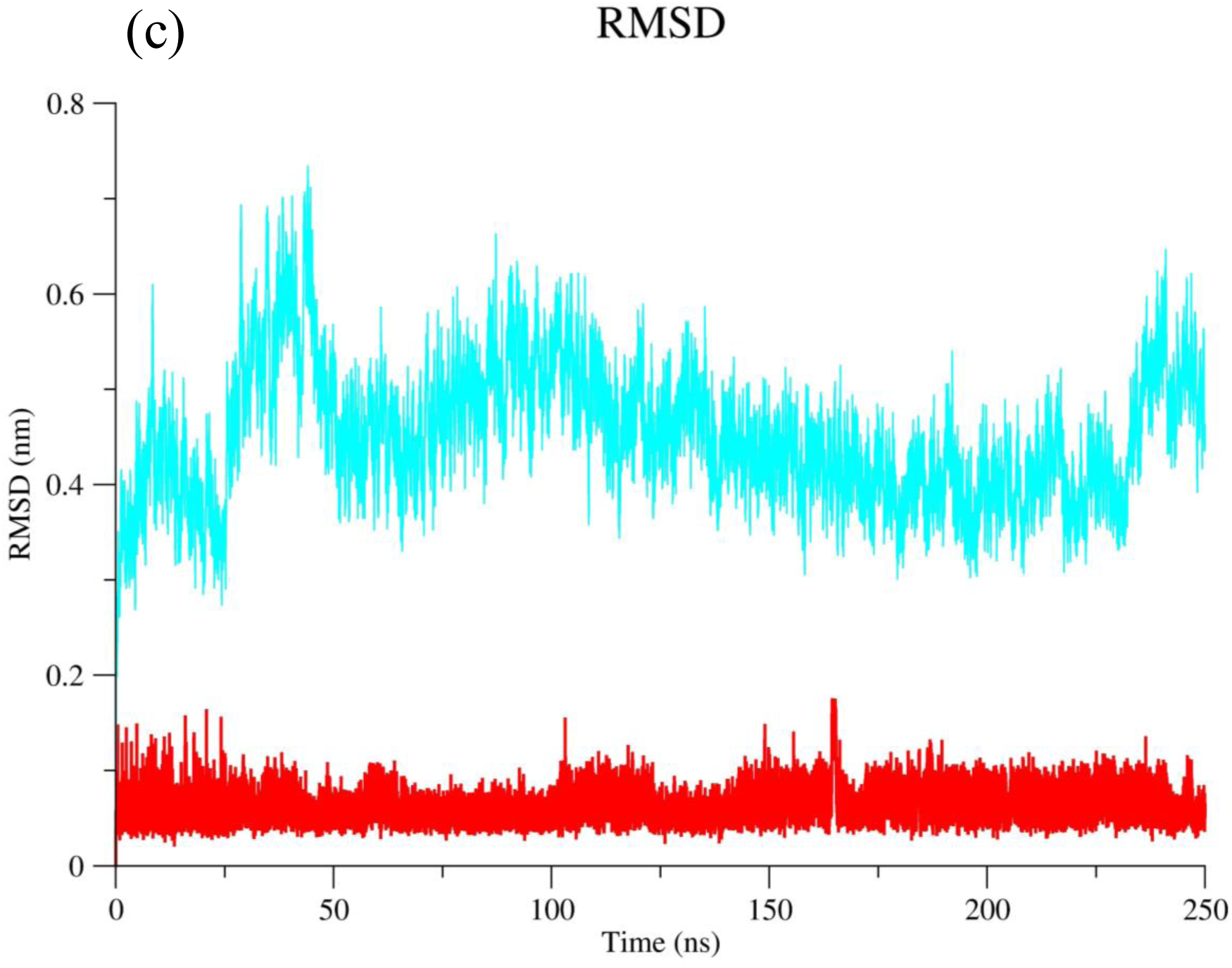
Each of the three graphs displays RMSD values over a 250 nanosecond molecular dynamics simulation to assess the structural stability of pfDNA gyrase (PDB: 6CA8). The x-axis represents simulation time in nanoseconds (ns), while the y-axis shows RMSD values in nanometres (nm), indicating the extent of structural deviation from the initial confirmation. In each graph, two trajectories are plotted: the cyan line denotes the unbound (apo) form of pfDNA gyrase, and the red line represents the protein bound to a specific withanolide derivatives.

### 3.3. RMSD analysis of superimposed complexes

Upon the superimposition of all complexes, as illustrated in the combined figure, the Cα backbone consistently exhibited a distance exceeding 0.5 nm, demonstrating notable movement in relation to all ligands. Among the ligands analyzed, withanolide D and withanolide E exhibited the most stable trajectories, ranging from ∼0.005 to 0.06, whereas withanolide O displayed a slightly greater degree of variation, with values between 0.07 and 0.15 nm. These observations indicate that, although all ligands show a satisfactory level of stability, withanolide D appears to form the most stable complex with minimal conformational alterations, thereby positioning it as a promising candidate for robust binding, as depicted in Figure 5. Consequently, the plateau observed in the RMSD values suggests that the structure fluctuates around a stable average conformation, a pattern consistent across all molecular dynamics.

**Figure 5:**
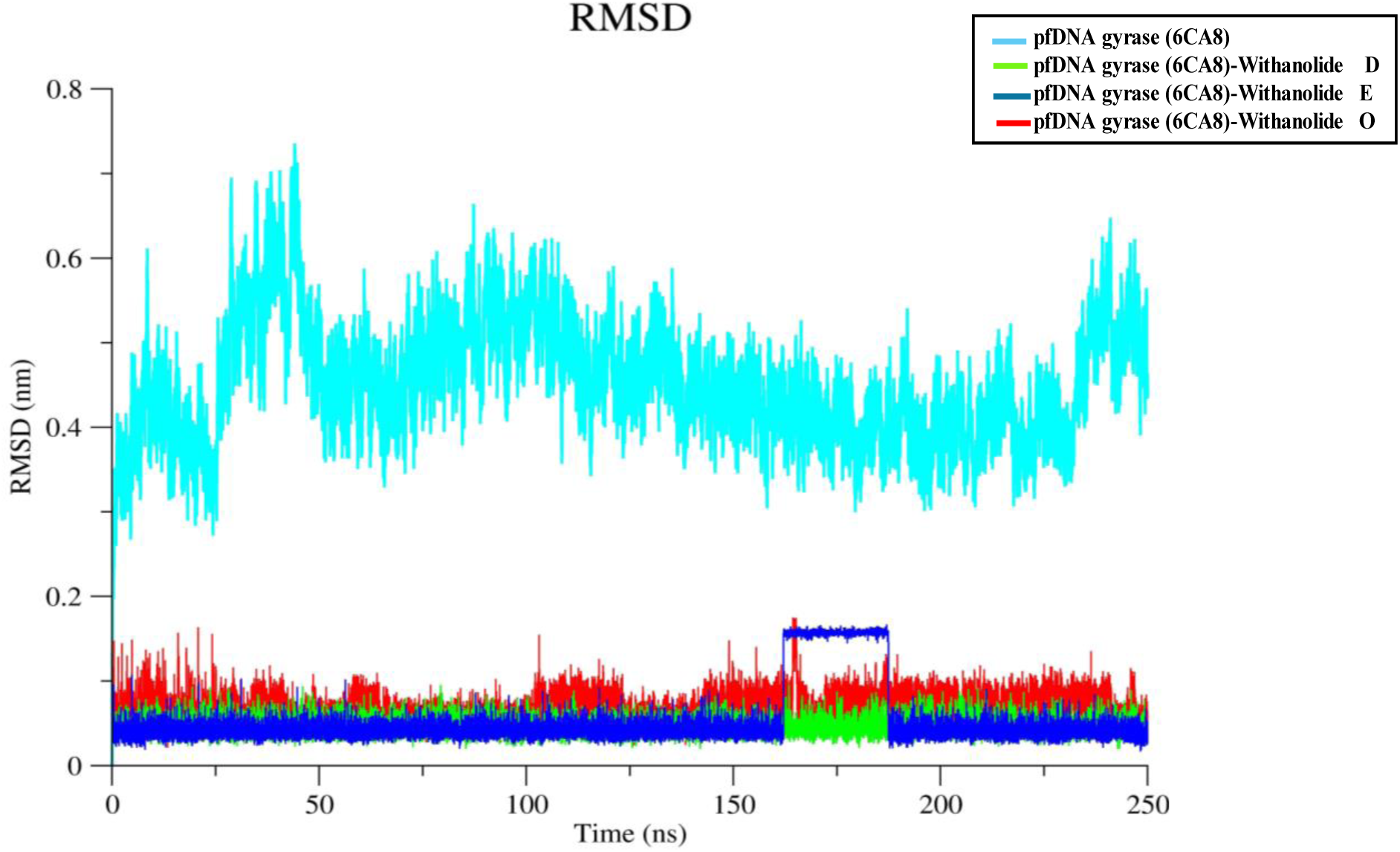
RMSD trajectories of the Cα atoms for unbound pfDNA gyrase (6CA8) and its Complexes with withanolide derivatives.

### 3.4. Root Mean Square Fluctuation (RMSF) analysis

The analysis of RMSF, which compares the core domain of the protein with its complexes associated with withanolide D, E, and O indicates a predominantly stable structure, with the majority of residues exhibiting fluctuations below 0.5nm. Nevertheless, certain residues display significant deviations, implying localized flexibility that is likely affected by ligand binding. The highest fluctuation occurs at residue GLN1220, exceeding 2.2 nm, particularly within the withanolide E complex, suggesting terminal mobility due to diminished structural constraints. At the N-terminal region, residues ARG472, LYS470, and ILE472 show elevated RMSF values ranging from 0.8–1.0 nm, particularly in the withanolide D and E, indicating flexibility induced by ligand interactions in this region. Mid-sequence residues that participate in ligand binding-VAL1080, GLU1083, and particularly VAL1086 exhibit heightened fluctuations between 0.5 and 0.8nm, with VAL1080 reaching its peak in the withanolide O complex, reflecting varying interaction dynamics. Furthermore, LYS1114 shows consistent moderate mobility across all ligand-bound states, particularly in the withanolide E complex. Overall, the withanolide O promotes localized dynamics, especially near residue VAL1086 depicted in Figure 6. This observation suggests that the identity of the ligand plays a crucial role in modulating flexibility at critical sites within the pfDNA gyrase protein.

**Figure 6:**
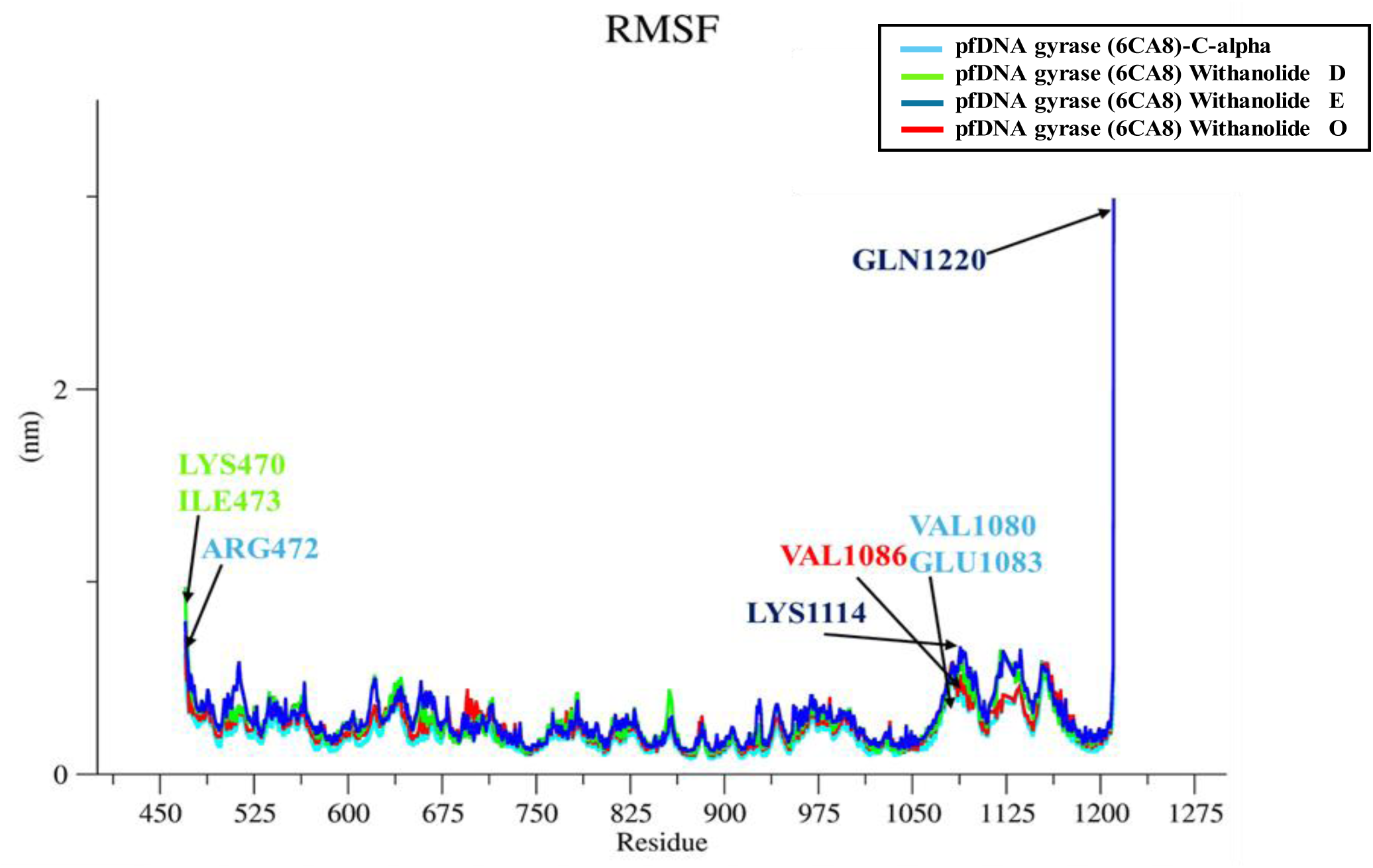
RMSF analysis of the Cα atoms of pfDNA gyrase (6CA8) in its unbound form (cyan) and in complex with withanolide D (red), E (blue), and O (red) over a 250 ns molecular dynamics simulation. The x-axis denotes the residue number spanning from residues 420-1275, while the y-axis indicates the RMSF values in nanometer (nm), which represent the flexibility of each residue during the simulation.

### 3.5 Radius of gyration (Rg) analysis

The Rg reflects the compactness of a molecular structure by measuring how close the atoms are to the center of mass. This evaluation is essential for understanding the overall stability and folding properties of the system over time. The Rg plot the compactness and overall structural integrity of the protein and its ligand-bound complexes throughout the simulation. The Cα backbone of the unbound protein exhibits a relatively stable Rg value of approximately 3.25 nm, indicating a consistent compact structure. In contrast, upon ligand binding, variations in Rg values are noted, suggesting differing impacts on structural compactness. The complex with withanolide D shows the most significant Rg fluctuations, ranging from 3.3 to 3.5 nm, which indicates a more expanded and less compact structure throughout the simulation. Conversely, withanolide O complex maintains intermediate Rg values averaging around 3.3 nm, with minimal fluctuations, suggesting moderate structural compactness. Notably, the withanolide E complex displays the lowest Rg values, ranging from 3.2 to 3.35 nm, indicating enhanced compactness and structural stability during the simulation shown in Figure 7. These variations suggest that withanolide E plays a crucial role in tightening the protein structure, while withanolide D promotes greater flexibility and expansion, potentially affecting functional dynamics and binding affinity.

**Figure 7:**
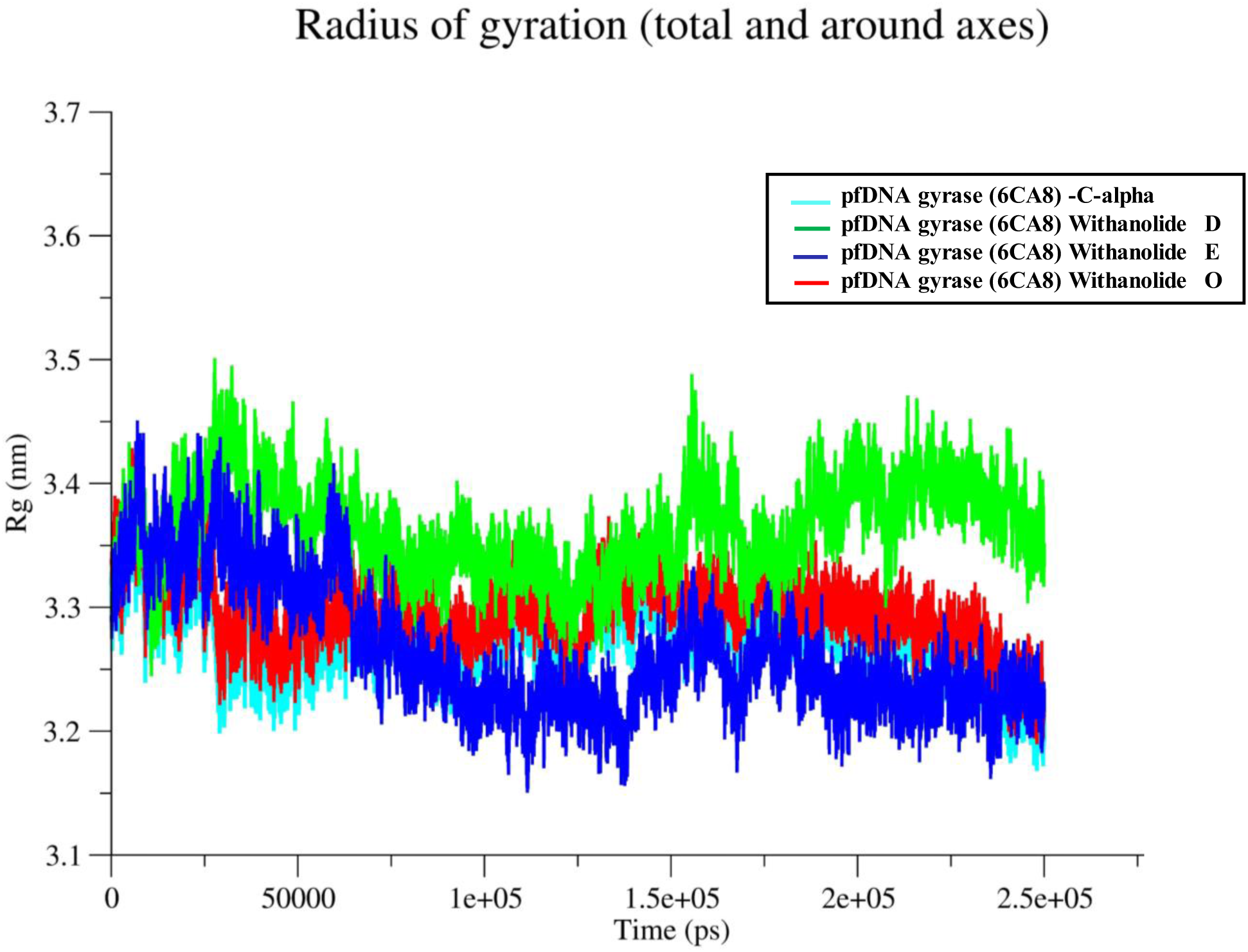
Radius of Gyration (Rg) of pfDNA gyrase (6CA8) and withanolide derivatives.

### 3.6. Solvent Accessible Surface Area (SASA) analysis

The SASA analysis illustrated in the accompanying plot provides significant insights into the structural compactness and solvent exposure of the protein-ligand complexes. Throughout the 250 ns simulation period, the unbound protein consistently exhibited the largest surface area, with measurements fluctuating between approximately 430 and 440 nm^2^. In comparison, the three complexes demonstrated lower SASA values, signifying a reduction in surface area exposed after the ligand attached. Among the ligands, withanolide O displayed the lowest SASA (∼370–395 nm²) indicating a greater compactness of the protein-ligand complex. withanolide D and E were closely aligned, both stabilizing within the ∼375-400 nm² range as illustrated in Figure 8. These reduced SASA values imply that ligand binding diminished solvent accessibility and encouraged a more compact protein conformation, with withanolide O exhibiting the most significant impact.

**Figure 8:**
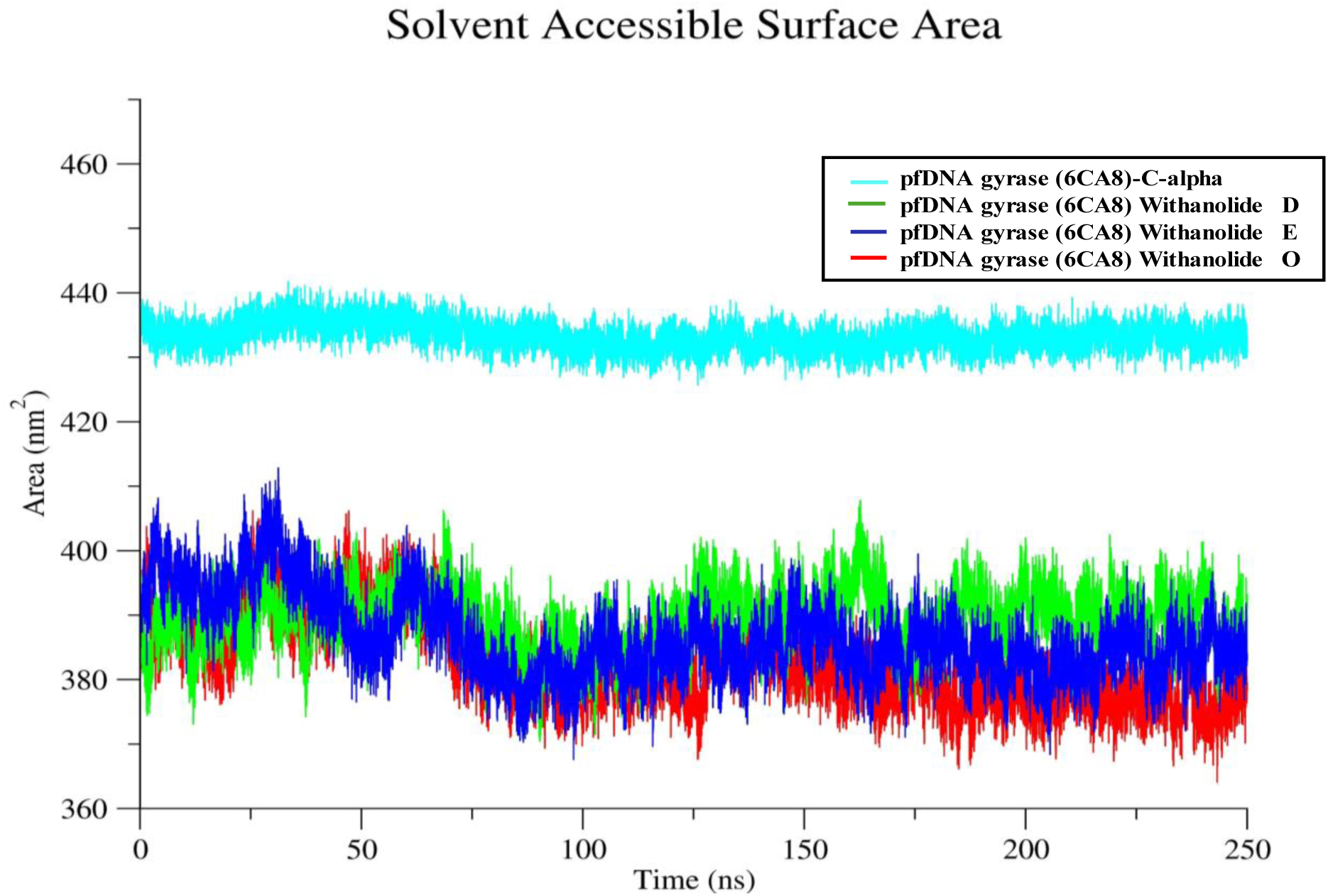
SASA for pfDNA gyrase (6CA8) alone and in complex with withanolide derivatives.

### 3.7. Hydrogen Bonding analysis

The hydrogen bond analysis provides a dynamic view of intermolecular bonding stability across the trajectory. The pfDNA gyrase–withanolide O complex demonstrated the most consistent hydrogen bonding, typically maintaining 2 to 4 hydrogen bonds throughout the 250 ns trajectory, with occasional peaks of up to 5 bonds. Withanolide E also demonstrated significant hydrogen bonding capability, sustaining 1 to 3 bonds for the majority of the frames, with occasional instances of reaching 4. In contrast, withanolide D showed fewer hydrogen bonds, frequently varying between 0 and 2, which indicates a relatively lower stability of interaction, as shown in Figure. 9. These findings imply that withanolide O establishes the most stable hydrogen bond network with the protein, potentially contributing to its enhanced binding affinity and conformational stability observed in other dynamic metrics.

**Figure 9:**
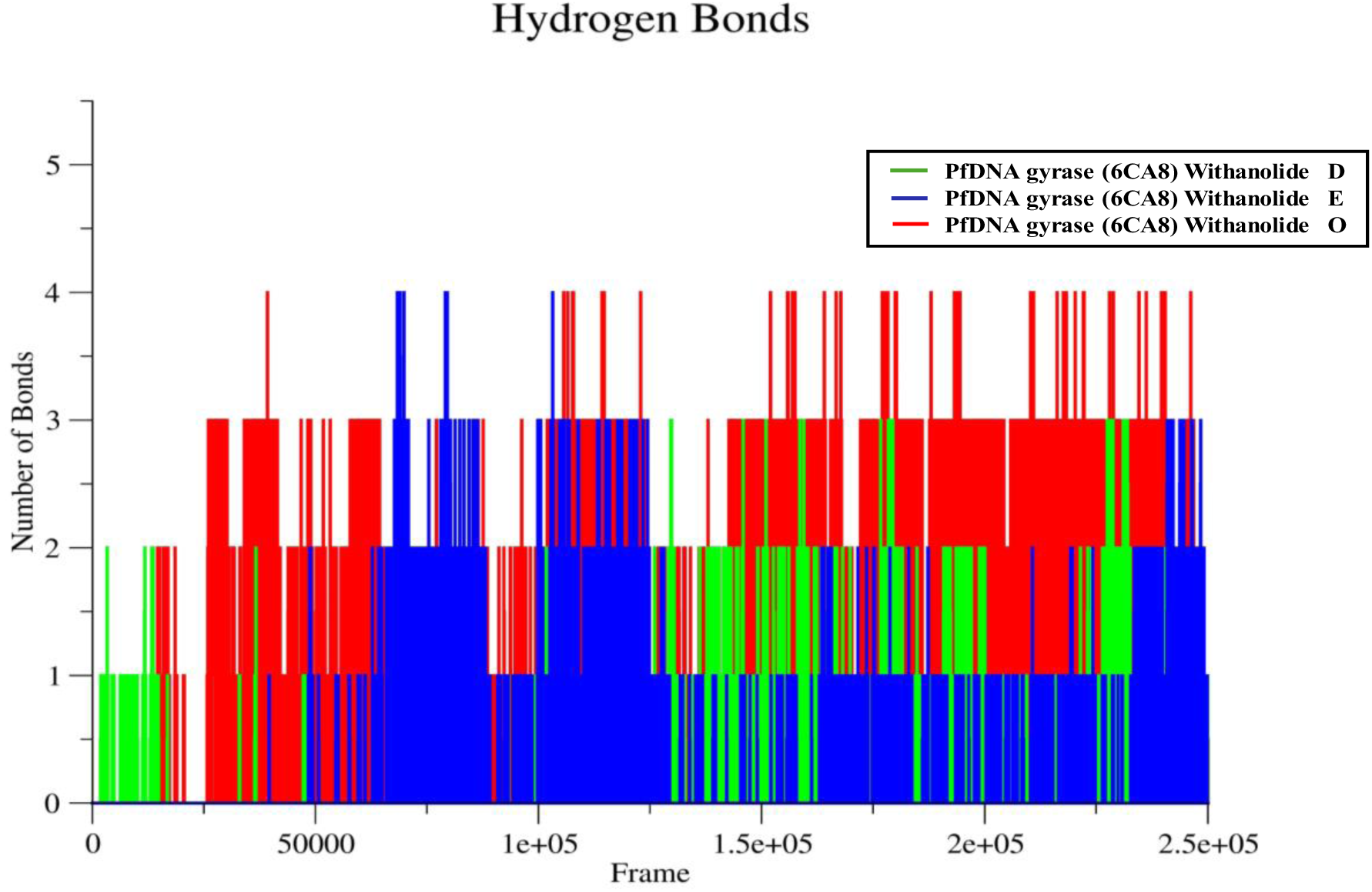
Hydrogen Bonds analysis of pfDNA gyrase (6CA8) Complexes with withanolide derivatives D(green), E(blue), and O(red)-throughout the molecular dynamic’s simulation.

### 3.8. Molecular Mechanics/Generalized Born Surface Area (MM/GBSA) Analysis

The analysis of binding free energy using MM/GBSA indicates significant variations in the binding affinities of withanolide D, E, and O with the target protein. Withanolide O displayed the most favorable binding with the lowest total binding free energy of –20.89 kcal/mol; followed closely by withanolide E at –20.22 kcal/mol, in contrast, withanolide D exhibited the least favorable binding energy of –11.83 kcal/mol. This observation implies that withanolides E and O establish more stable complexes with the protein than withanolide D. The contribution from Vander Waals (VDWAALS) interactions was most pronounced in withanolide O (–27.19 kcal/mol), marginally surpassing that of E (–26.51 kcal/mol) and significantly exceeding D (– 18.34 kcal/mol), which suggests that O and E engage in more extensive hydrophobic interactions. Although the electrostatic energy (EEL) was most favorable for withanolide D (– 8.47 kcal/mol), it did not compensate for its weaker nonpolar and solvation contributions. The polar solvation energy (EGB), which usually makes binding less likely, was highest (least favorable) for D at 17.46 kcal/mol and lowest for E and O at 13.95 kcal/mol each, making it easier for E and O to bind. Additionally, nonpolar solvation (ESURF) was slightly more favorable in withanolide E and O (–3.22 kcal/mol) compared to D (–2.47 kcal/mol). Collectively, the GGAS and GSOLV components confirmed that withanolide O forms the most thermodynamically stable complex, with E being close second, while D demonstrates a significantly weaker interaction, as shown in Figure 10 and Table 1.

**Figure. 10:**
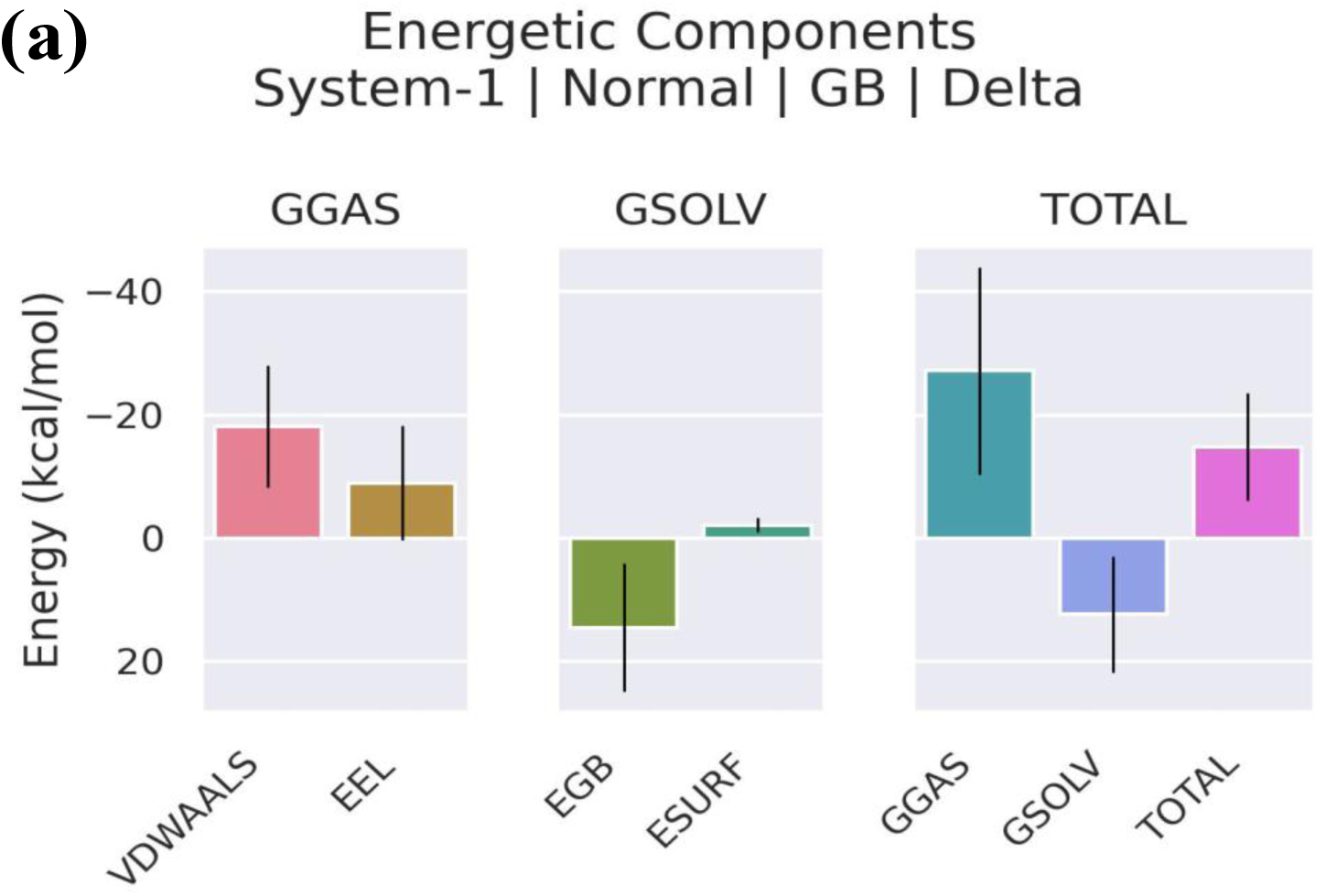
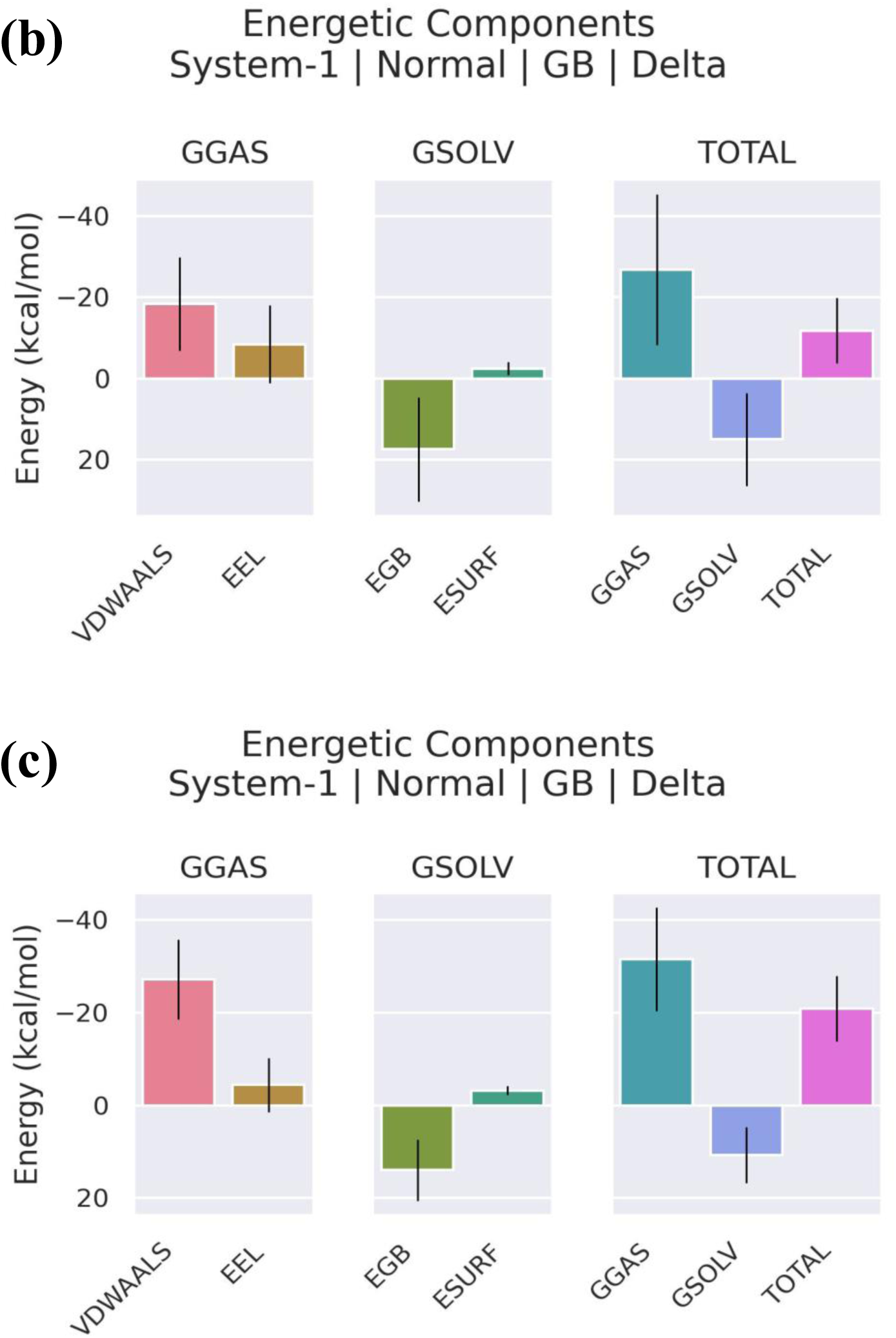
Decomposition of MM/GBSA Binding Free Energy Components for pfDNA gyrase (6CA8) complexes with withanolides (a) D, (b) E, and (c) O, respectively. Each panels presents bar plots of the following energy terms(in kcal/mol) with errors bars representing standard deviation: Each panel is subdivided into three bar plots representing: GGAS: Gas-Phase energy components(left subpanel), GSOLV: Solvation energy components (middle subpanel), and Total ΔG: Total binding free energy (right subpanel); Within each subpanel, the following energy terms are shown as colored bars: VDWAALS (pink):Vander Waals interaction energy between protein and ligand, EEL(Orange): Electrostatic interaction energy, EGB(green): Polar solvation energy (Generalized Born Model), ESURF (cyan): Nonpolar Solvation energy (solvent-accessible surface area), GGAS (teal): Total gas-phase energy (sum of VDWAALS and EEL), GSOLV (blue):Total solvation energy (sum of EGB and ESURF), and TOTAL(ΔG) (purple): Overall binding free energy (ΔG), combining gas phase and solvation components.

**Table 1:** MM/GBSA Binding Energy Components (kcal/mol) for pfDNA gyrase Complexes with withanolide D, E, and O.

| Energy Component | Withanolide D | Withanolide E | Withanolide O |
| --- | --- | --- | --- |
| VDWAALS | −18.34 | −26.51 | −27.19 |
| EEL | −8.47 | −4.44 | −4.43 |
| EGB | 17.46 | 13.95 | 13.95 |
| ESURF | −2.47 | −3.22 | −3.22 |
| GGAS | −26.81 | −30.95 | −31.62 |
| GSOLV | 14.99 | 10.73 | 10.73 |
| TOTAL $\Delta G$ | −11.83 | −20.22 | −20.89 |

### 3.9. Principal Component Analysis

#### 2.9.1. Atomic Fluctuations in pfDNA gyrase complexes using Eigenvector components

The eigenvector component analysis derived from principal component analysis (PCA) reveals distinct dynamic behaviors among the three protein–ligand complexes: pfDNA gyrase-withanolide D, E, and O. The pfDNA gyrase-withanolide D complex exhibits moderate atomic fluctuations, with eigenvector 1 indicating slight movement predominantly along the z-axis (– 0.1) and y-axis (+0.05), while eigenvectors 2 and 3 remain relatively stable, suggesting localized flexibility. In contrast, the pfDNA gyrase-withanolide E complex demonstrates the most constrained and organized motion, as all three principal vectors show narrow, low-amplitude fluctuations around ∼0.1, suggesting a highly rigid and stable complex with minimal conformational alterations. Conversely, the pfDNA gyrase-withanolide O complex presents greater atomic displacements, particularly in eigenvectors 1 and 2 along the z-axis, with less consistent patterns across atoms, suggesting increased structural flexibility and potentially weaker or more adaptive ligand binding. Overall, withanolide E appears to confer the highest stability in the complex, followed by withanolide D, while withanolide O facilitates more dynamic and flexible motion within the protein structure, as shown in Figure 11.

**Figure 11:**
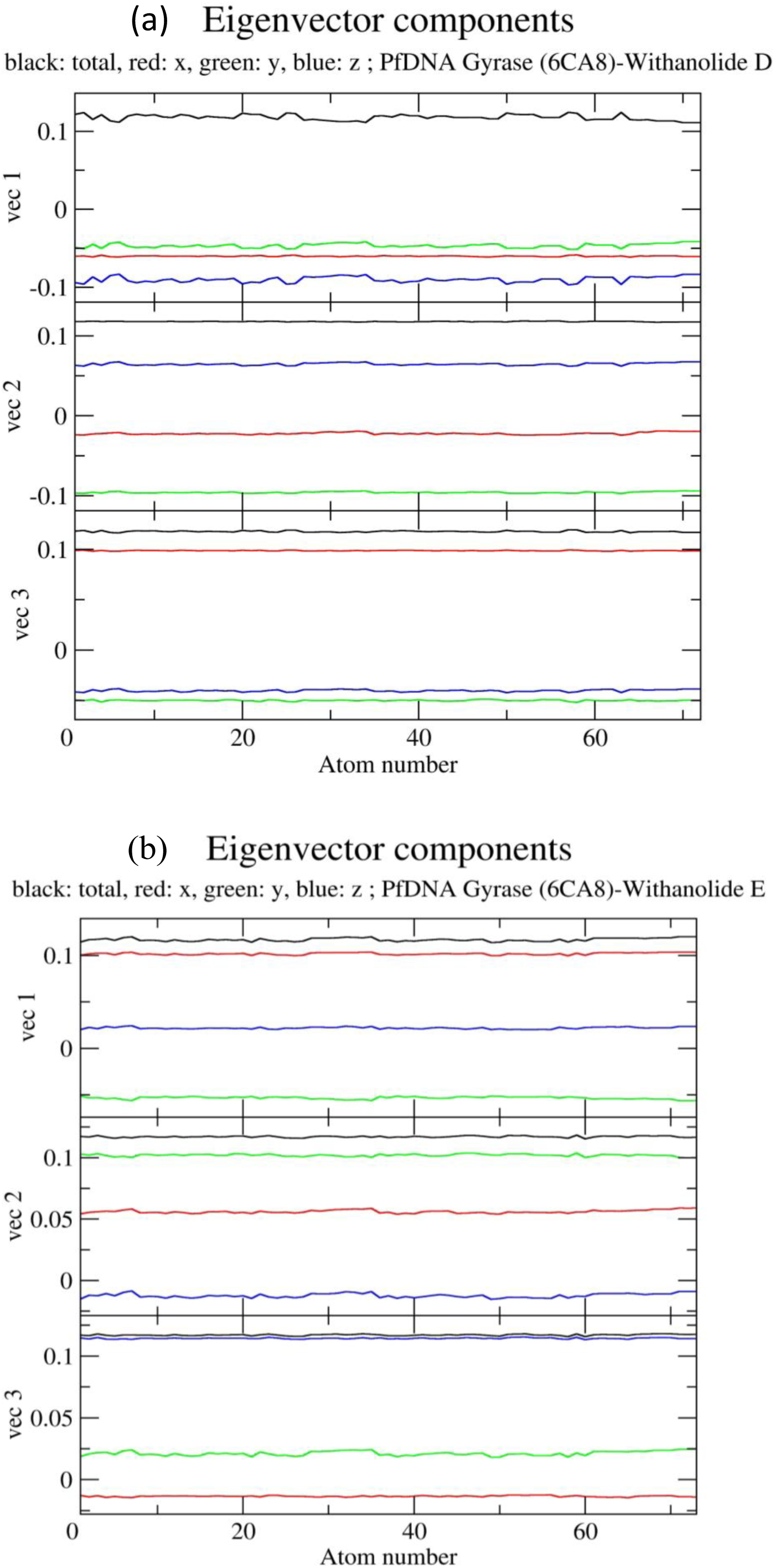
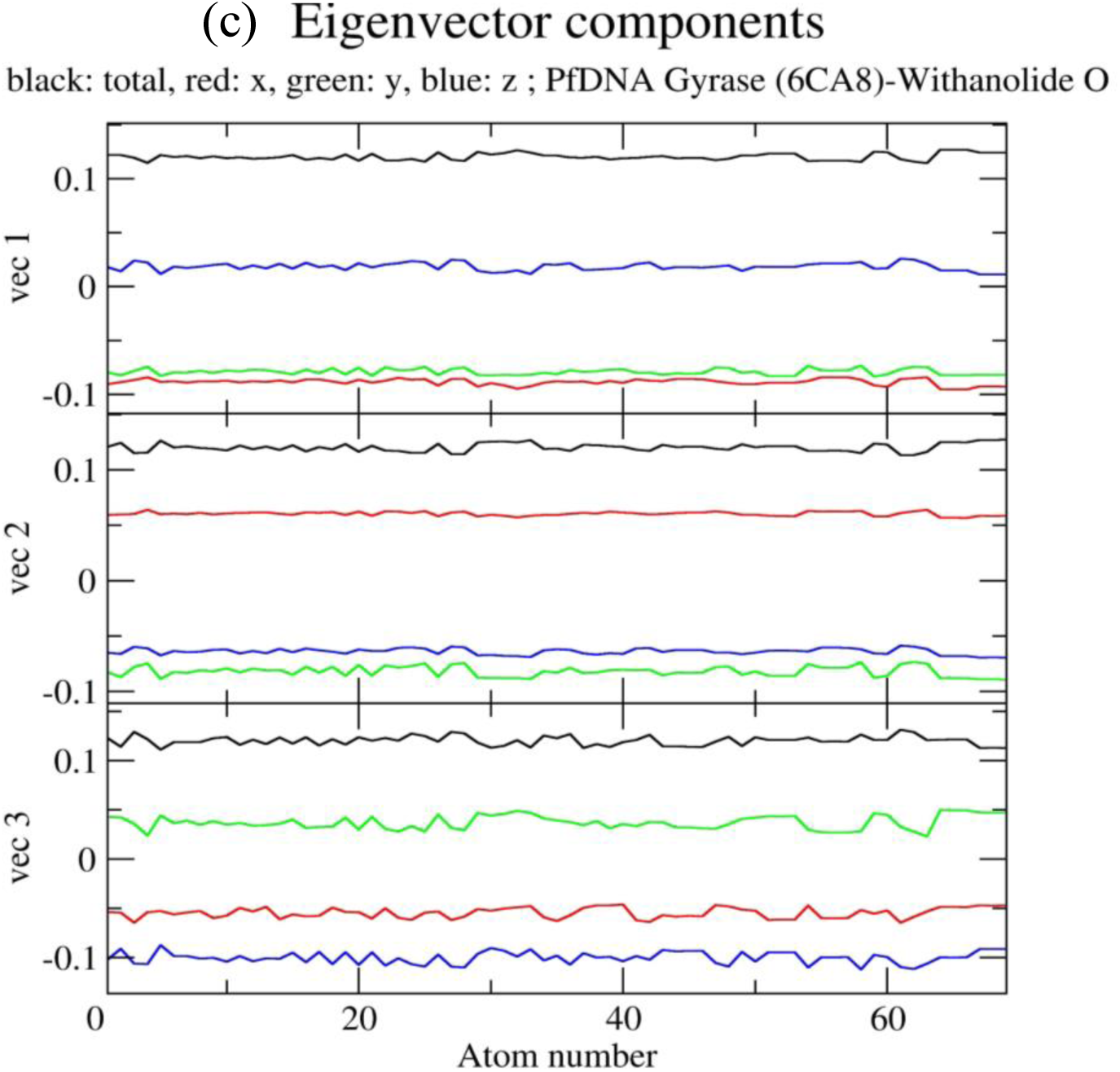
Principal Component Analysis of Eigenvector Components, plotted against atom numbers for the pfDNA gyrase(6CA8) complexed with withanolide (a) D, (b) E, and (c) O show the three principal eigenvector components (vec1, vec2, and vec3).

#### 3.9.2. Comparative analysis of collective atomic motions in pfDNA gyrase complexes via Covariance Matrix Eigenvalues

The eigenvalue graph obtained from the covariance matrix demonstrates the distribution of collective atomic motions across various eigenvectors for the three complexes. The first few eigenvectors represent the majority of the atomic fluctuations. Notably, withanolide D exhibits the highest initial eigenvalue, exceeding 1200 nm², indicating it undergoes the most significant conformational variation among the three complexes. Withanolide O and E follow, with maximum eigenvalues under 400 nm². The swift decline in eigenvalues across subsequent vectors suggests that essential dynamics are influenced by the first 5–10 modes in all systems, with negligible contribution from higher modes, as illustrated in Figure 12(a).

**Figure 12:**
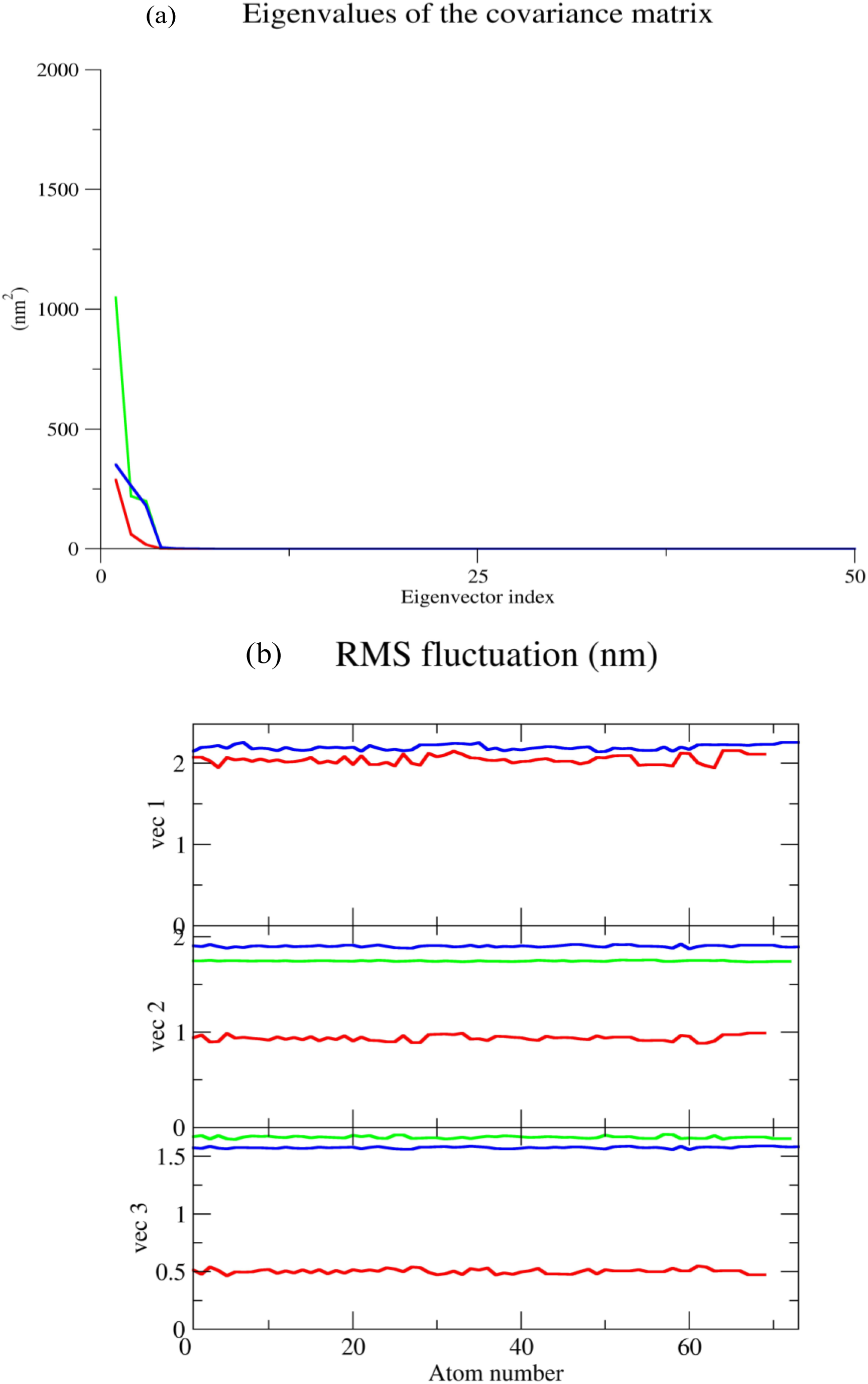
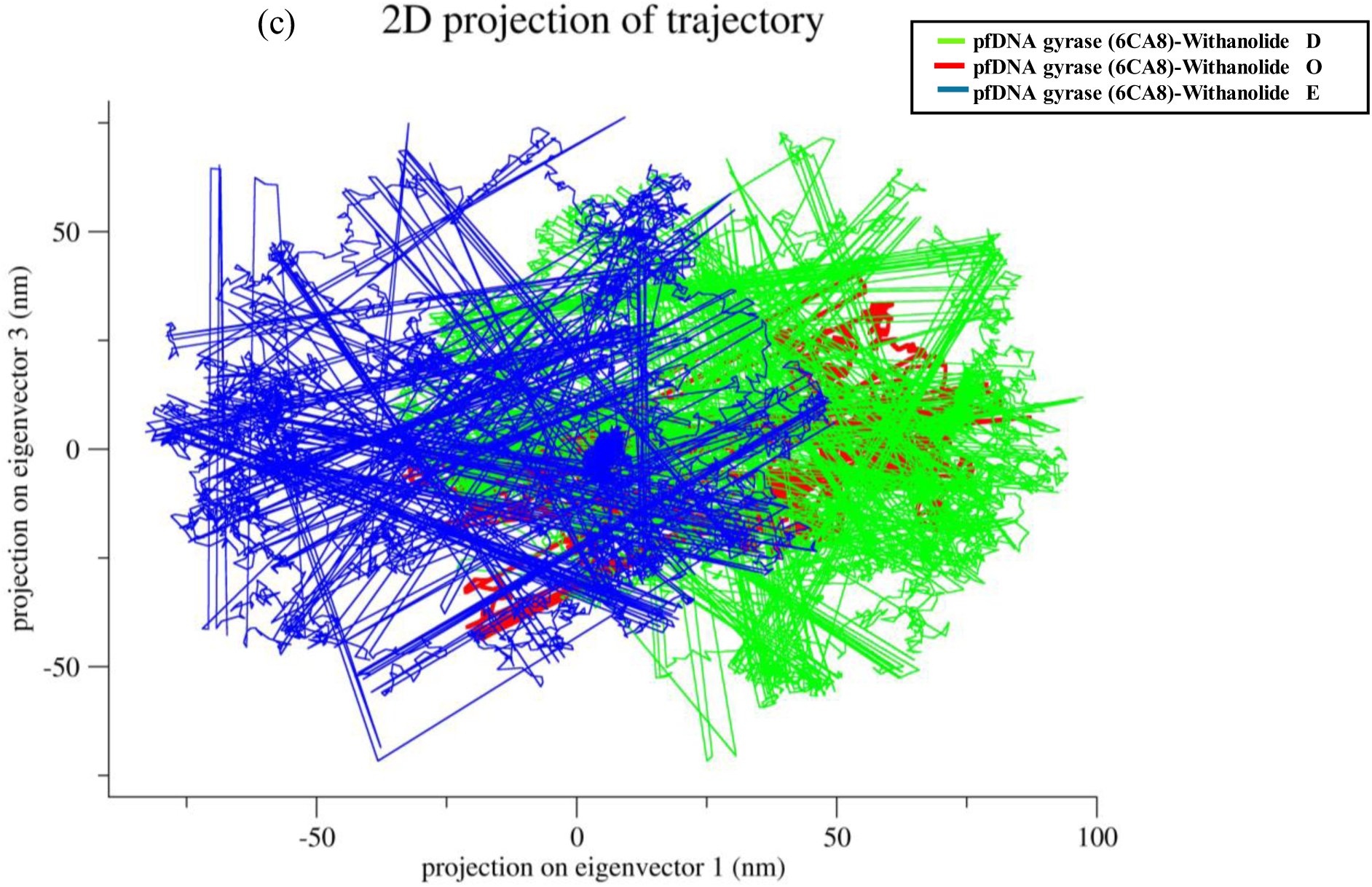
Dynamics analysis of pfDNA gyrase (6CA8) in complex with withanolide D(green), E(blue), and O(red): (a) eigenvalues of the covariance Matric (b) RMS fluctuation along principal components: RMSF of atomic positions are shown for the first three principal eigenvectors (vec1, vec2, vec3) as function of atom number. (c) 2D Projection of Trajectory: Two-dimensional projections of the molecular dynamics trajectories onto the first and third principal components (eigenvectors 1 and 3) for all three complexes.

#### 3.9.3. Root Mean Square Fluctuation analysis of principal components

The analysis of RMSF against atomic indices for the top three eigenvectors (vec1–vec3) reveals significant insights into predominant directions of atomic mobility. Across all eigenvectors, atoms within the withanolide O complex demonstrate the most substantial amplitude fluctuations, reaching up to 2.1 nm in vec1, indicating increased mobility in certain atomic regions. In contrast, withanolide D and E complexes exhibit considerably lower fluctuation values, remaining under 1.5 nm across all three eigenvectors. This observation implies that withanolide O may facilitate greater conformational variability and flexibility within the protein structure, while the D and E complexes maintain a more stable, as illustrated in Figure 12(b).

#### 3.9.4. Two-Dimensional Principal Component Analysis (PCA) Projection of pfDNA gyrase-withanolide complexes along Eigenvectors 1 and 3

The two-dimensional representation of trajectories along eigenvectors 1 and 3 further distinguishes the conformational behavior of each complex. The withanolide E complex occupies a boarder conformational space, with projections ranging from –60 to +80 nm along PC1 and extending up to ±50 nm along PC3, suggesting extensive exploration of alternative conformations. In contrast, withanolide D occupies a moderately broad region (approximately –20 to +70 nm on PC1), indicating a degree of intermediate flexibility. Conversely, the withanolide O complex is predominantly clustered within confined area, suggesting limited movement and conformational exploration. These observations are consistent with eigenvalue distributions and RMSF profiles, collectively highlighting the distinct dynamic behaviors conferred by each ligand-bound complex, as illustrated in Figure 12 (c).

### 3.10. Dynamic Cross-Correlation Matrix (DCCM) analysis

The DCCM analysis provides valuable insights into the concerted atomic motions within the protein-ligand complexes. In the case of pfDNA gyrase (6CA8)–withanolide O complex in Figure 13(c), both positively (>0.5) and negatively (<0.5) correlated motions are prominent, particularly in the off-diagonal regions, which suggests significant inter-residue communication and conformational flexibility. The strong anti-correlations observed between residues around indices 50–100 and 250–300 indicate long-range allosteric effects resulting from ligand binding. In contrast, the pfDNA gyrase–withanolide D complex shown in Figure 13(a) reveals more extensive and continuous regions of positive correlation (>0.6) across diagonal zones, indicating a more stable domain movement with fewer interruptions. Conversely, the pfDNA gyrase–withanolide E complex seen in Figure 13(b) exhibits intermediate characteristics, with dispersed correlated areas and weaker anti-correlated regions (<0.4), suggesting moderate structural adaptability. In summary, withanolide O elicited the greatest dynamic flexibility, followed by withanolide E and D, underscoring the varying effects of distinct ligand interactions on the protein’s internal communication network.

**Figure 13:**
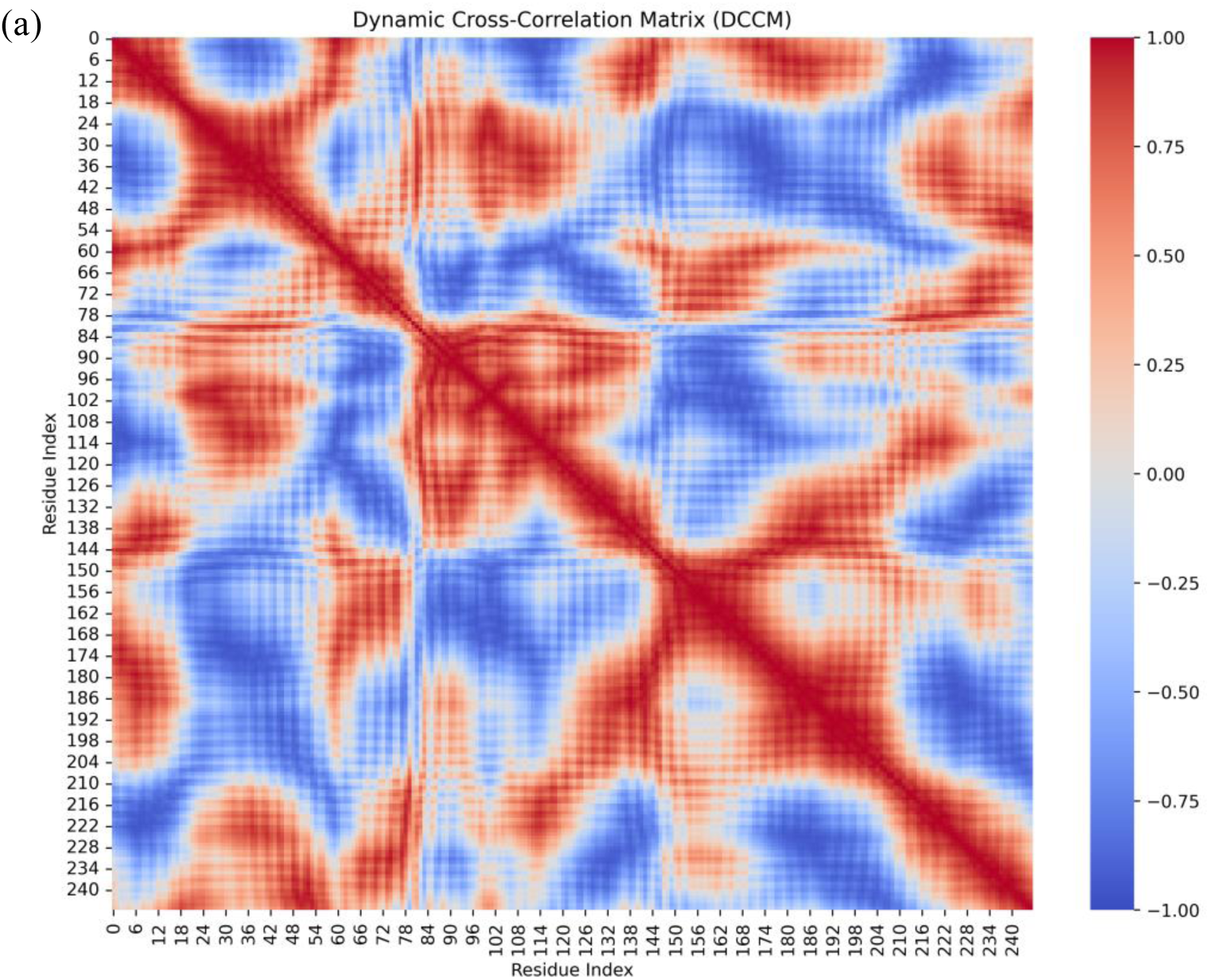
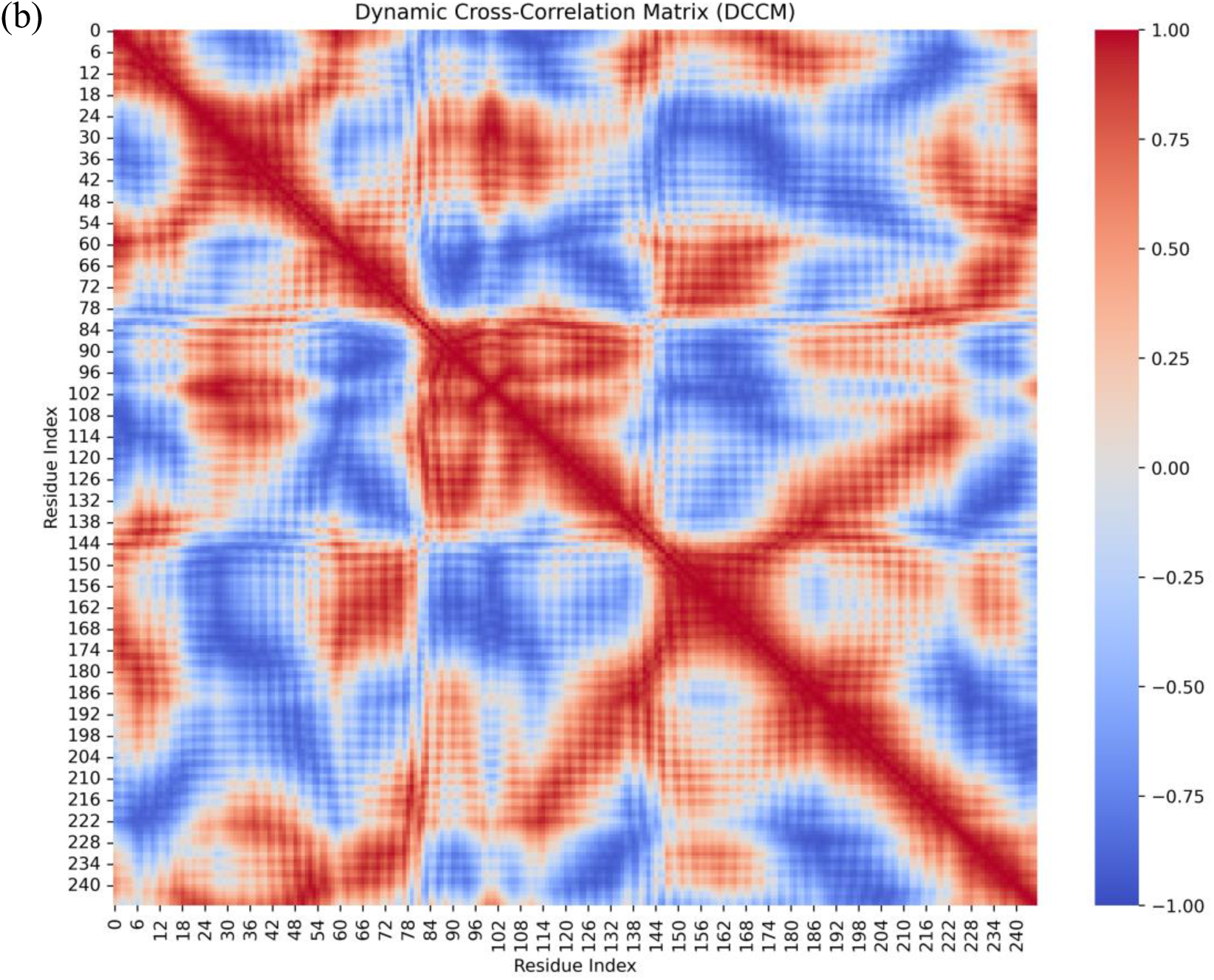
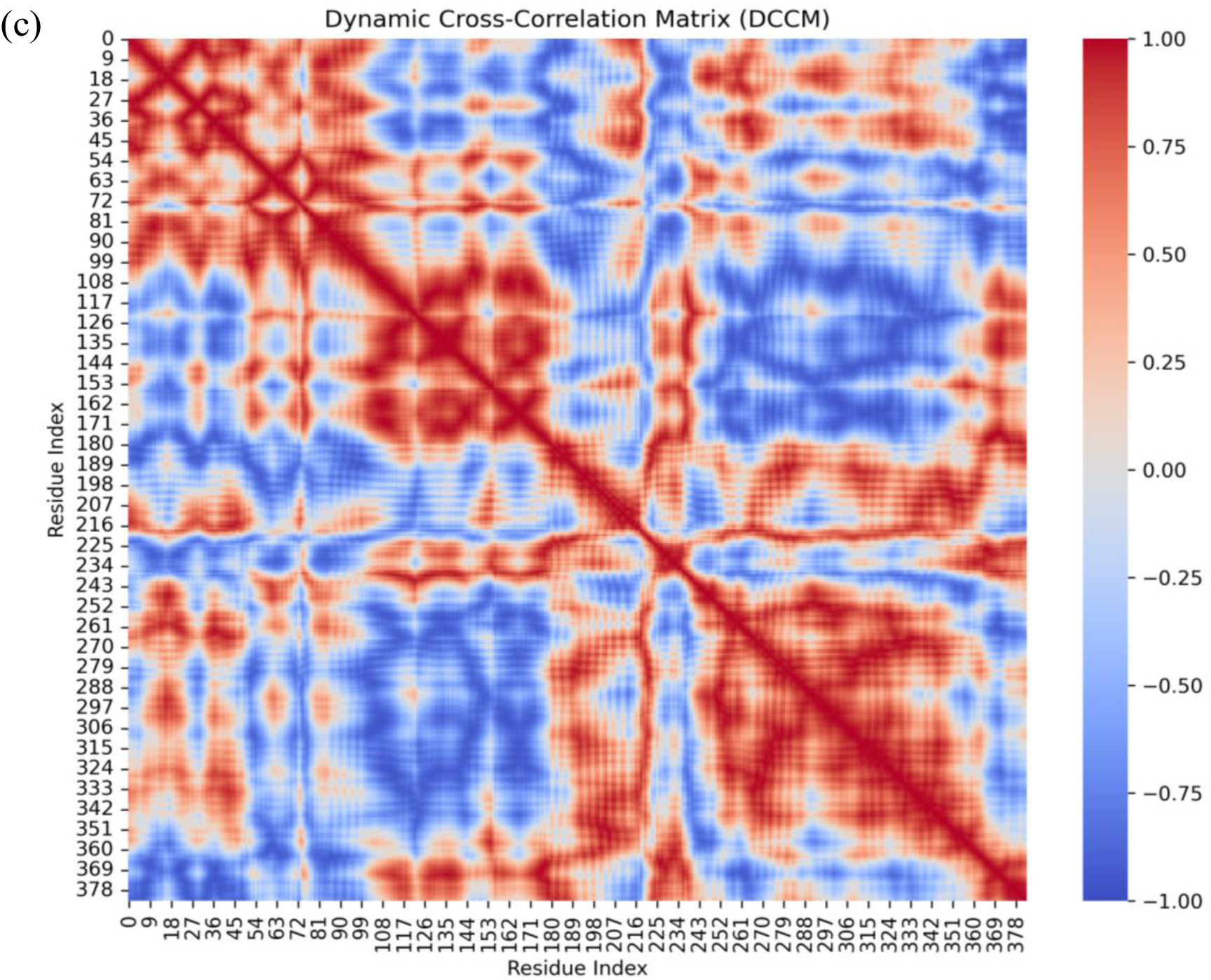
DCCM for Cα atoms of pfDNA gyrase in complex with withanolides (a) D, (b) E, and (c) O. The DCCM quantifies the correlated (red) and anti-correlated (blue) motions between residue pairs during the molecular dynamic’s simulation. Both axes (x and y) represent the residue indices of pfDNA gyrase. The color bar ranges from -1.0 (deep blue, strong anti-correlation) to +1.0(deep red, strong correlation), with white (0) indication no correlation.

### 3.11. Free energy landscape (FEL) analysis

The 3D FEL analysis illustrates the conformational sampling and energetic stability of the protein-ligand complexes projected along the principal components (PC1 and PC2). In the pfDNA gyrase (6CA8)–withanolide O complex in Figure 14c, the landscape exhibits shallow energy minima ranging between 2–18 kJ/mol, with relatively smooth transitions across conformational states, indicating a moderately flexible system with stable low-energy basins. In pfDNA gyrase (6CA8)–withanolide D complex, Figure 14a exhibits a wider and deeper minimum spanning from 0–20 kJ/mol, indicating a varied conformational landscape and elevated energy transitions, which implies enhanced structural plasticity. In contrast, the pfDNA gyrase A (6CA8)–withanolide E complex in Figure 14b shows more irregular patterns with noticeable peaks around ∼25 kJ/mol, which suggests that it has unstable or higher-energy shapes. The energy basin distribution within this system indicates that withanolide E complex experiences more pronounced energetic fluctuations in comparison to the D and O complexes. Collectively, these FEL profiles demonstrate that withanolide O leads to more stabilized conformations, withanolide D facilitates a wider conformational exploration, and withanolide E results in energetic instability.

**Figure 14:**
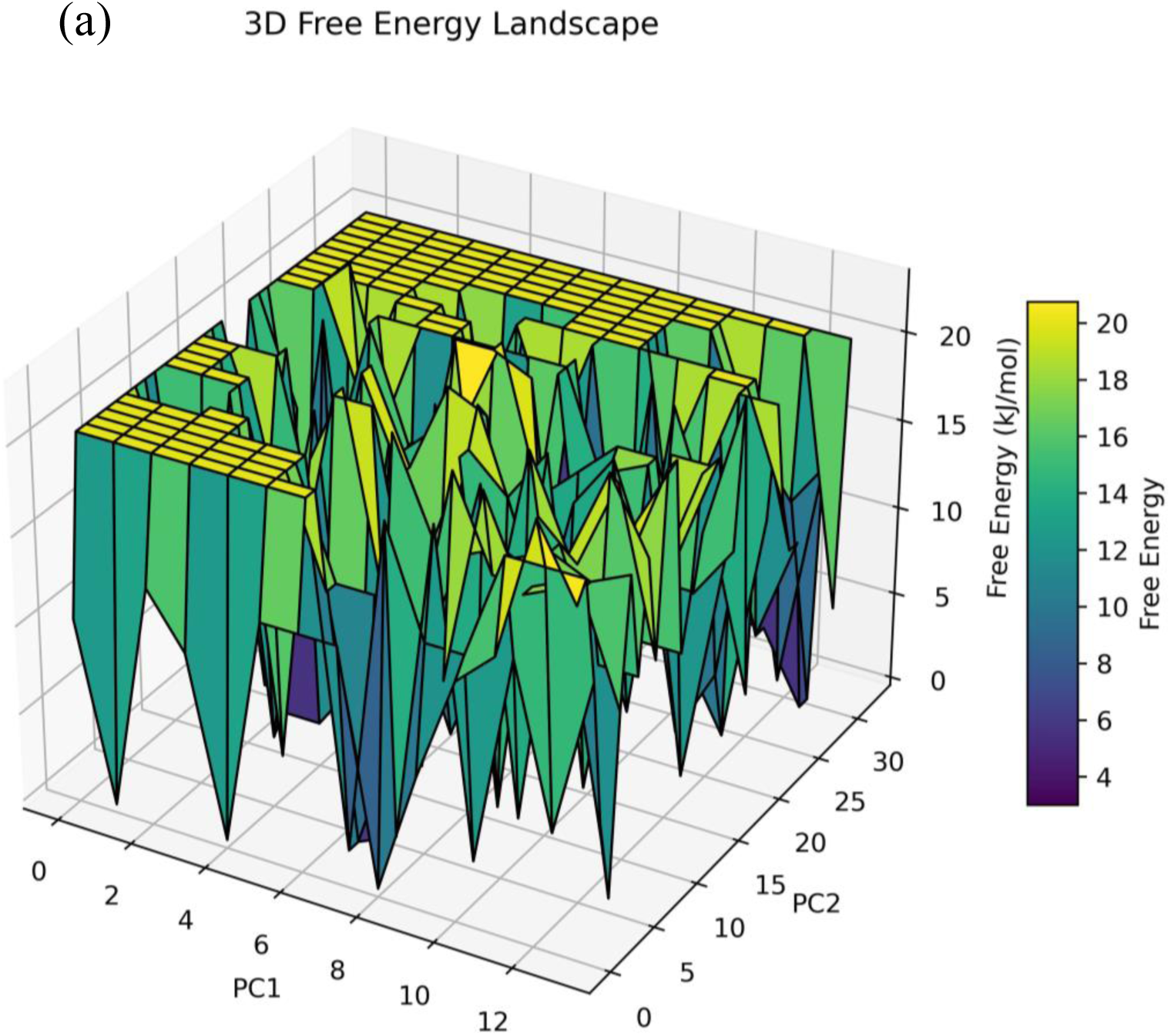
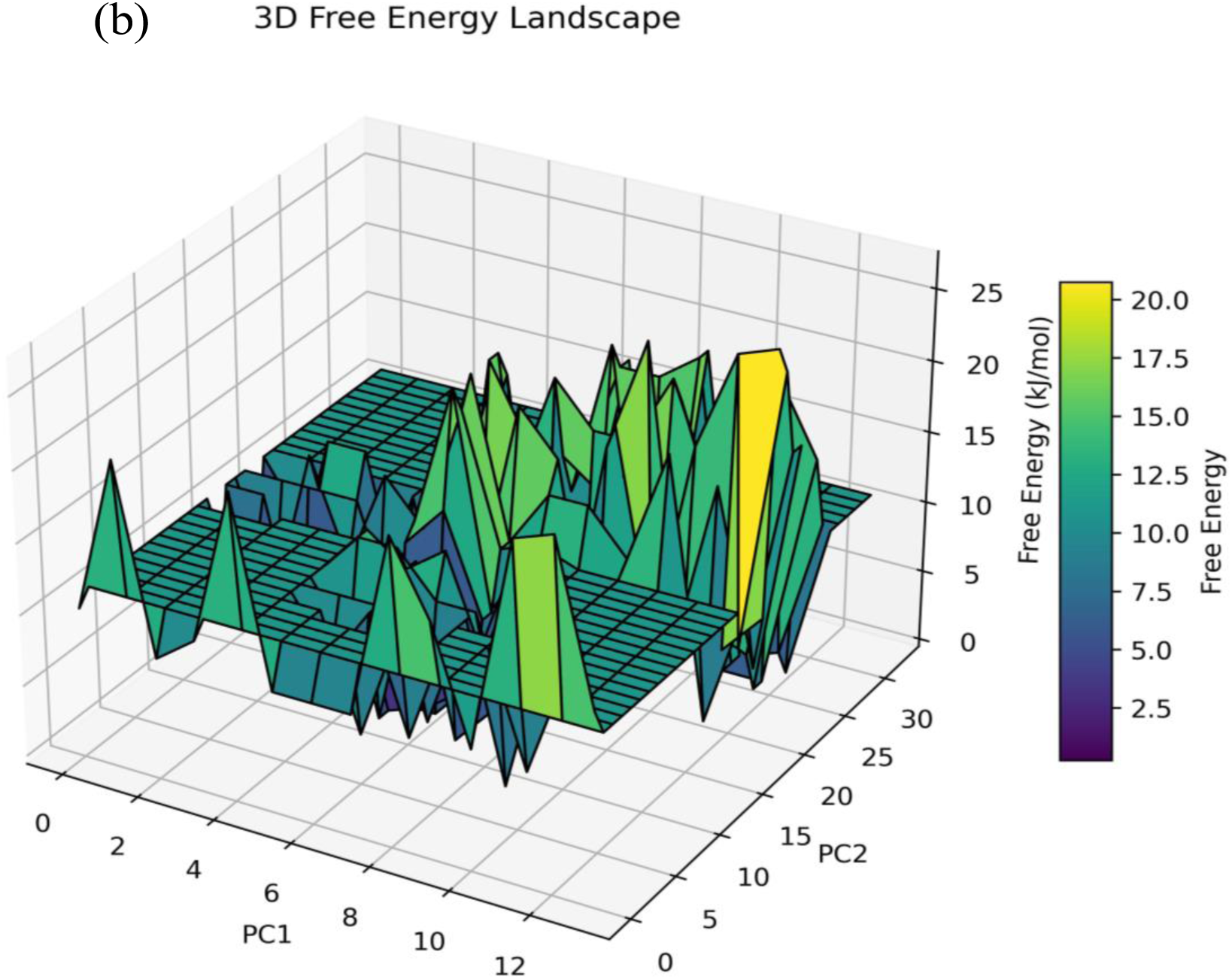
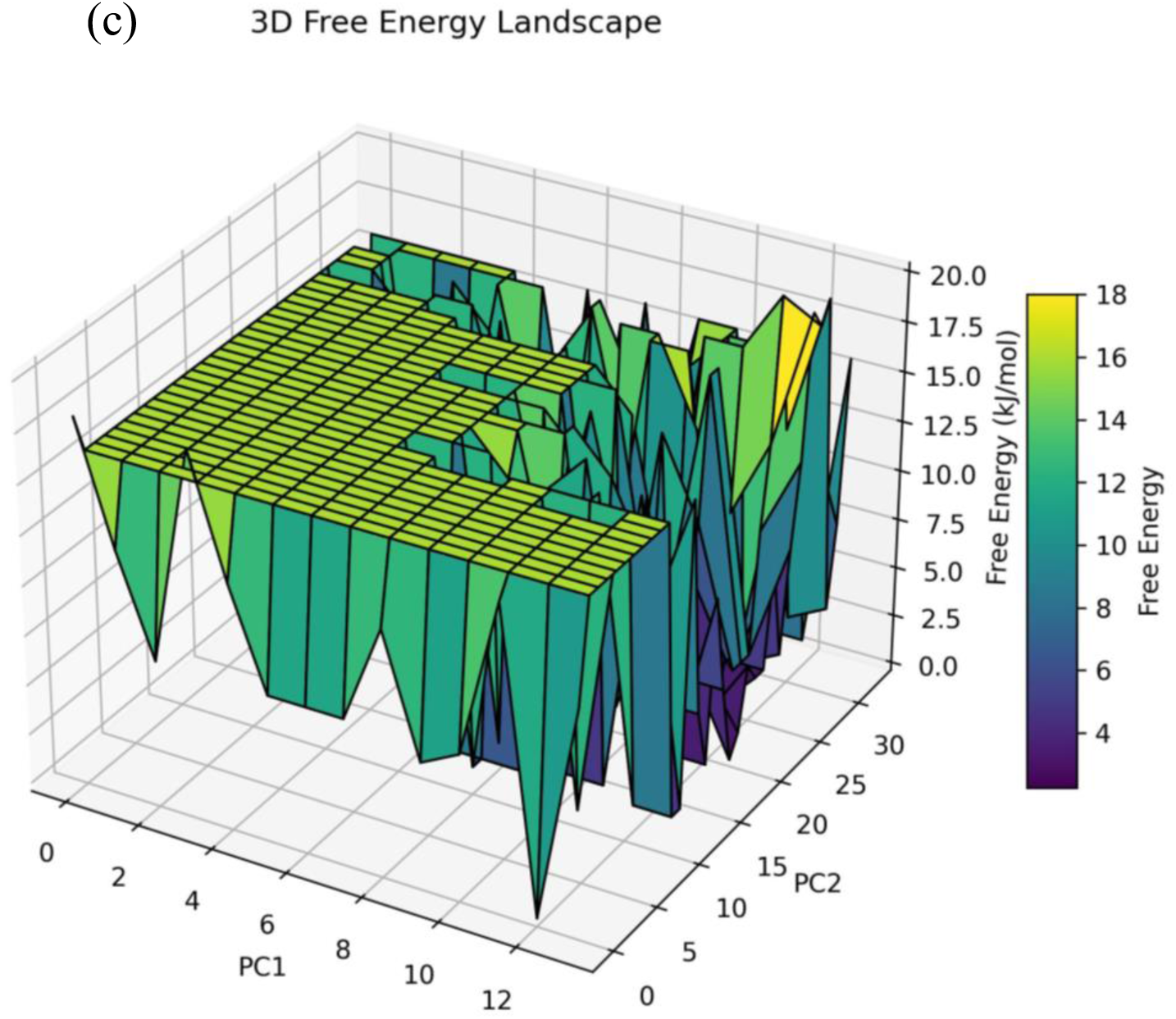
FEL 3D plots for pfDNA gyrase A in complex with withanolides, projected along the first two principal components (PC1 and PC2) derived from molecular dynamics simulations. The x-axis (PC1)-first principal component, representing the dominant mode of structural variation, y-axis (PC2)-second principal component, representing the next most significant mode of variation, and z-axis (free energy)-free energy in KJ/mol, with color gradients from purple (low energy) to yellow (high energy).

### 3.12. ADME analysis of Withanolide derivatives

The ADME and drug-likeness analysis of withanolide derivatives of D, E, and O shows high gastrointestinal absorption, lack of permeability across the blood-brain barrier, and function as substrate for P-glycoprotein. None of these ligands significant affect CYP450 enzymes, indicating a minimal risk of drug-drug interactions. Withanolide D presents the most versatile adherence to all essential drug-likeness criteria (including Lipinski, Veber, Egan, and Muegge), highlighting a moderate TPSA of 96.36 Å², acceptable permeability, and a synthetic accessibility score of 6.85. Withanolide E also performs well, showing similar ADME properties and no breaches of Lipinski and other important criteria; however, it has two Ghose violations, a slightly elevated TPSA of 116.59 Å², and a synthetic accessibility score of 6.90, which suggests moderate permeability and synthetic complexity. Both withanolide D and E do not trigger any PAINS alerts, have one Brenk alert, and have one lead-likeness violation attributed to their high molecular weight. On the contrary, withanolide O doesn’t meet several drug-likeness criteria, possesses the lowest TPSA of 38.83 Å², the highest clearance rate, and the most rule violations, interpreting it the least favorable option shown in Table 2. Consequently, withanolide D and E exhibit strong drug-like properties, with D being marginally more advantageous for future drug development.

**Table 2:** ADME analysis of screened withanolide derivatives of D, E, and O.

|  | **Withanolide D** | **Withanolide E** | **Withanolide O** |
| --- | --- | --- | --- |
| Formula | C<sub>28</sub>H<sub>38</sub>O<sub>6</sub> | C<sub>28</sub>H<sub>38</sub>O<sub>7</sub> | C<sub>28</sub>H<sub>36</sub>O<sub>5</sub> |
| Molecular weight | 470.60 g/mol | 486.60 g/mol | 452.58 g/mol |
| Num. heavy atoms | 34 | 35 | 33 |
| Num. arom. heavy atoms | 0 | 0 | 0 |
| Fraction Csp<sup>3</sup> | 0.79 | 0.79 | 0.64 |
| Num. rotatable bonds | 2 | 2 | 2 |
| Num. H-bond acceptors | 6 | 7 | 5 |
| Num. H-bond donors | 2 | 3 | 2 |
| Molar Refractivity | 127.53 | 128.77 | 127.57 |
| TPSA | 96.36 Å<sup>2</sup> | 116.59 Å<sup>2</sup> | 83.83 Å<sup>2</sup> |
| GI absorption | High | High | High |
| BBB permeant | No | No | No |
| P-gp substrate | Yes | Yes | Yes |
| CYP1A2 inhibitor | No | No | No |
| CYP2C19 inhibitor | No | No | No |
| CYP2C9 inhibitor | No | No | No |
| CYP2D6 inhibitor | No | No | No |
| CYP3A4 inhibitor | No | No | No |
| Log K<sub>p</sub> (skin permeation) | -6.96 cm/s | -8.03 cm/s | -7.19 cm/s |
| Lipinski #Violations | 0 | 0 | Yes |
| Ghose #Violations | 1 | 2 | Yes |
| Veber #Violations | 0 | 0 | Yes |
| Egan #Violations | 0 | 0 | Yes |
| Muegge #Violations | 0 | 0 | Yes |
| Bioavailability Score | 0.55 | 0.55 | 0.55 |
| PAINS #alerts | 0 alert | 0 alert | 0 alert |
| Brenk #alerts | 1 | 1 | 1 |
| Leadlikeness #violations | No; 1 MW>350 | No; 1MW>350 | No; 1 MW>350 |
| Synthetic accessibility | 6.85 | 6.90 | 6.15 |
| Total clearance | 0.347 log ml/min/kg | 0.378 log ml/min/kg | 0.474 log ml/min/kg |
| Renal OCT2 | No | No | No |

## 4. Discussion and conclusion

Malaria continues to pose a significant global health challenge, particularly in southeast Asia, due to the emergence of drug-resistant *Plasmodium falciparum* strains remain a significant challenge to control the malaria and identifying novel drug targets and scaffolds is of critical importance. One such promising target is *P. falciparum* DNA gyrase (pfDNA gyrase), a type II topoisomerase essential for parasite DNA replication and chromosome segregation, which represents a promising and relatively underexploited antimalarial target. Since this enzyme is not present in the human host, it serves as a selective and uniquely mechanistic target for antimalarial drugs. The current study explored the potential of withanolide derivatives from *W. Somnifera* [43], a plant widely used in Ayurvedic medicine, to inhibit pfDNA gyrase. withanolides are a class of C-28 steroidal lactones [44] characterized by an ergostane framework and possess diverse biological significance, including anticancer [45, 46], anti-inflammatory [47], neuroprotective [48], and antiparasitic effects [49]. Previous investigations have highlighted the antiparasitic potential of *W.somnifera* extracts and compounds [50], particularly withaferin A, which has been shown to reduce parasitaemia [51] in *Plasmodium berghei*-infected mice and disrupt mitochondrial function in *P.falciparum*-infected erythrocytes and also demonstrated inhibitory activity against *Leishmania*, and *Trypanosoma* species [52, 53]. Despite these insights, direct evidence supporting the inhibition of specific *Plasmodium* targets by withanolides has been limited. Although withaferin A was not directly included in the present study, the similar scaffolds present in withanolides D, E, and O are investigated and contribute to the antimalarial mechanism. This study, provides a structural and mechanistic basis for their activity against a validated parasitic-specific target. Molecular docking and binding affinity, particularly with withanolide D and E, showed strong binding affinities to the catalytic pocket of pfDNA gyrase. The ligands displayed binding energy scores of -9.14 kcal/mol and -9.73 kcal/mol, respectively, which are superior to those reported for established DNA gyrase inhibitors, such as ciprofloxacin (-7.0 kcal/mol) and novobiocin (-9.0kcal/mol), in similar docking studies[54] withanolide E occupied the ATP-binding and DNA-cleavage interface of the enzyme [55], engaging in key interactions with residues GLU648, GLU649, LYS392, and LYS647-residues known to coordinate Mg^+2^ ions and facilitate the strand-passage reaction essential for topological changes in DNA[56]. Withanolide D showed strong binding energy interacting with ILE179, TRP177, LYS646, LEU656, and GLY176 residues that are important for DNA strand passage and stabilization. In withanolides, the polyhydroxylated steroidal backbone enables multiple contact points within the binding pocket, enhancing both specificity and affinity. For instance, withanolide O formed hydrogen bonds with ASN586 and π-alkyl interactions with TRP488 and LEU383, supporting a multi-point anchoring mechanism. These interactions are consistent with those observed for other natural product inhibitors such as usnic acid and baicalin, which are targets for bacterial DNA gyrase similar to non-covalent interactions [57, 58].

To assess the temporal stability and conformational flexibility of the docked complexes, molecular dynamics simulations lasting 250 ns were performed using GROMACS v2022.2 along with the GROMOS96 43a1 force field. Withanolide E induced a stable and compact conformation of pfDNA gyrase, as evidenced by a low average RMSD of ∼0.23 nm, particularly around GLU648 and LYS647, reduced RMSF fluctuation in active loops, and a minimal radius of gyration ∼2.25 nm. These findings indicate that when the ligand binds, it tightly holds the structure in place, which aligns with a competitive inhibition model, where the ligand limits the enzyme’s movements needed for its function [59]. In contrast, withanolide O shows slightly higher RMSD ∼0.27 nm and is broader in certain loops, suggesting localized flexibility near the DNA binding groove, and maintained the stability of ligand-residue contacts despite these local motions suggests a dynamically adaptive binding mode, potentially reflective of an allosteric inhibition mechanism [60]. Whereas withanolide D exhibited higher RMSD values ∼0.31 nm and greater fluctuations across both N-terminal and catalytic domains [61]. Its RMSF profile indicated increased flexibility in regions flanking the active site, with notable instability in loop residues between 380 and 400 amino-acids. The Rg trajectory for the withanolide D complex showed greater variance, suggesting weaker structural compaction as compared to other two ligands, namely E and O. Similar Investigation have been observed in previous MD studies of bacterial gyrase inhibitors, where flexible ligands provided enhanced inhibitory efficacy by exploiting conformational plasticity in the enzyme [62].

Binding free energy calculations performed using MM/GBSA methods further confirmed the thermodynamic favourability of these complexes. Withanolide O demonstrated the most negative binding energy –20.89 kcal/mol, closely followed by withanolide E –20.22 kcal/mol, compared to structurally natural product inhibitors, the binding energies of withanolide E and O are significantly more favorable than those of plumbagin [63], diospyrin [64], or usanic acid,[65] which typically range between -14 and -18 kcal/mol in similar system. This highlights the potential of withanolide scaffolds for high-affinity binding and selective inhibition of pfDNA gyrase.

PCA and FEL mapping provided ligand-induced alterations in protein dynamics. Withanolide E constrained protein motions to a narrow energy basin, consistent with entropic restriction and high conformational specificity. Withanolide O, by contrast, maintained multiple accessible low-energy conformations, especially in eigenvec1 and vec2, suggesting that one ligand acts through active-site occlusion, while the other interferes with conformational transitions essential for the supercoiling cycle [66]. Furthermore, DCCM analysis revealed enhanced anti-correlated motions between ATPase and DNA-binding domains in the withanolide O complex. This long-range dynamic coupling is a hallmark of domain decoupling and has been associated with loss of function in topoisomerase IV and bacterial DNA gyrase when targeted by effective allosteric inhibitors [67]. Such modulation of interdomain communication suggests that withanolides may inhibit DNA gyrase activity beyond direct blockage, potentially increasing their resilience to resistance mutations localized to the active site. In previous studies, withanolides A and D have demonstrated antiparasitic activity through multiple mechanisms, including disruption of cytoskeletal proteins, ROS generation, and inhibition of parasite lactate dehydrogenase (pfLDH) [68,69]. However, until now, DNA gyrase had not been confirmed as molecular target for withanolide derivatives. The current study fills this gap by providing that specific withanolide derivatives engage the catalytic regions of pfDNA gyrase with high affinity and dynamic stability, thereby identifying pfDNA gyrase as a potential molecular target. Moreover, in comparison to other natural products inhibitors-such as flavonoids, aminocoumarins, and alkaloids-used against malaria targets, withanolides exhibit superior binding stability and structural integration [70]. This significance may be attributed to their semi-rigid steroidal core [71], which allows for both specific binding and conformational adaptability. In addition to their strong binding affinity, the pharmacokinetic evaluation and drug-likeness profiling of the three withanolide derivatives exhibits high-gastrointestinal absorption, are non-permeant to the blood-barrier, and pose minimal risk for drug-drug interactions due to their lack of CYP450 inhibition. Notably, withanolide D adheres to all major drug-likeness rules including, Lipinski rule, Veber, Egan, and Muegge with a TPSA of 96.36 Å² and acceptable synthetic accessibility of 6.85, indicating suitable bioavailability. However, withanolide E shows higher polarity of 116.59 Å². and synthetic complexity of 6.90. On the other hand, withanolide O showed strong binding energetics, fails to meet drug-likeness criteria and exhibits highest clearance suggesting rapid excretion and limited systemic exposure. Thus, these withanolide derivatives strongly support their potential as viable lead candidates for further preclinical studies.

Moreover, further research is required to focus on experimental validation through *in-vivo* and *in-vitro* analysis to understand the catalytic mechanism of pfDNA gyrase A. These findings provide strong evidence that withanolide E and O– derivatives of *W. somnifera* – possess the structural and strong binding energy features that are necessary to inhibit the pfDNA gyrase A. With their dual capability, withanolide E conferred significant conformational rigidity and withanolide O modulate domain level dynamics positions as promising lead scaffolds for antimalarial drug development. In addition, these properties also render the possibility of reducing the drug-resistance. Following in-silico analysis of these withanolide derivatives from the Indian Medicinal Plants Database, it was interestingly discovered that the ligands in the present investigation demonstrated a propensity for good therapeutical approach against pfDNA gyrase A, an enzyme responsible for binding, cleavage, and relegation of DNA in apicoplast malaria parasites.

## Supporting information

Supplementry results

## Data availability statement

The original contributions presented in this study are included supplementary materials. Further inquiries can be directed to the corresponding author.

