## Supplementry results for "Mechanistic Insights into the inhibition of *Plasmodium falciparum* DNA gyrase A by withanolides derivatives through integrated computational analysis"

Supplementary results

```
Program: needle
Rundate: Sat 12 APRIL 2025 20:36:26
Commandline: needle
-auto
-stdout
-asequence emboss_needle-I20250517-203616-0752-60176758-p1m.asequence
-bsequence emboss_needle-I20250517-203616-0752-60176758-p1m.bsequence
-datafile EBLOSUM62
-gapopen 10.0
-gapextend 0.5
-endopen 10.0
-endextend 0.5
-aformat3 pair
-sprotein1
-sprotein2 Align_format: pair Report_file: stdout Aligned_sequences: 2
1: EMBOSS_001
2: EMBOSS_001
Matrix: EBLOSUM62 Gap_penalty: 10.0
Extend_penalty: 0.5
Length: 751
Identity:	731/751 (97.3%)
Similarity:	731/751 (97.3%) Gaps:		20/751 ( 2.7%)
Score: 3799.0
pfDNA gyrase Sequence ======================================= EMBOSS_001	1 --RERIIGIPKLEDANDAGSKYSQECTLILTEGDSAKTSCLAGLSIVGRD
48
```

| EMBOSS_001 | 1 | \|\|\|\|\|\|\|\|\|\|\|\|\|\|\|\|\|\|\|\|\|\|\|\|\|\|\|\|\|\|\|\|\|\|\|\|\|\|\|\|\|\|\|\|\|\|\|\|  MARERIIGIPKLEDANDAGSKYSQECTLILTEGDSAKTSCLAGLSIVGRD |
| --- | --- | --- |
| 50  EMBOSS_001 | 49 | KYGVFPLKGKLLNVRDASFKQLMDNKEIQNIFRIMGLDITDKNKDDIKGL |
| 98 |  | \|\|\|\|\|\|\|\|\|\|\|\|\|\|\|\|\|\|\|\|\|\|\|\|\|\|\|\|\|\|\|\|\|\|\|\|\|\|\|\|\|\|\|\|\|\|\|\|\|\| |
| EMBOSS_001 100 | 51 | KYGVFPLKGKLLNVRDASFKQLMDNKEIQNIFRIMGLDITDKNKDDIKGL |
| EMBOSS_001 148 | 99 | RYGSLMIMTDQDYDGSHIKGLLINMIHKFWPSLLKHKGFLSEFVTPIVKV  \|\|\|\|\|\|\|\|\|\|\|\|\|\|\|\|\|\|\|\|\|\|\|\|\|\|\|\|\|\|\|\|\|\|\|\|\|\|\|\|\|\|\|\|\|\|\|\|\|\| |

| EMBOSS_001 | 101 | RYGSLMIMTDQDYDGSHIKGLLINMIHKFWPSLLKHKGFLSEFVTPIVKV |
| --- | --- | --- |
| 150 |  |  |
| EMBOSS_001 | 149 | QKGSQEYSFFTIAEYEQWKENTNLLGWKIKYYKGLGTSTDREFKQYFSDI |
| 198 |  |  |
|  |  | \|\|\|\|\|\|\|\|\|\|\|\|\|\|\|\|\|\|\|\|\|\|\|\|\|\|\|\|\|\|\|\|\|\|\|\|\|\|\|\|\|\|\|\|\|\|\|\|\|\| |
| EMBOSS_001 | 151 | QKGSQEYSFFTIAEYEQWKENTNLLGWKIKYYKGLGTSTDREFKQYFSDI |
| 200 |  |  |
| EMBOSS_001 | 199 | KNHKIMFLWTGDRDGDSIDMAFSKKRIEDRKLWLQNFILGSYVDHKEKDL |
| 248 |  |  |
|  |  | \|\|\|\|\|\|\|\|\|\|\|\|\|\|\|\|\|\|\|\|\|\|\|\|\|\|\|\|\|\|\|\|\|\|\|\|\|\|\|\|\|\|\|\|\|\|\|\|\|\| |
| EMBOSS_001 | 201 | KNHKIMFLWTGDRDGDSIDMAFSKKRIEDRKLWLQNFILGSYVDHKEKDL |
| 250 |  |  |
| EMBOSS_001 | 249 | SYYDFVNKELIYYSRYDTERSIPNIMDGWKPGQRKVLYGCFKRNLRNECK |
| 298 |  |  |
|  |  | \|\|\|\|\|\|\|\|\|\|\|\|\|\|\|\|\|\|\|\|\|\|\|\|\|\|\|\|\|\|\|\|\|\|\|\|\|\|\|\|\|\|\|\|\|\|\|\|\|\| |
| EMBOSS_001 | 251 | SYYDFVNKELIYYSRYDTERSIPNIMDGWKPGQRKVLYGCFKRNLRNECK |
| 300 |  |  |
| EMBOSS_001 | 299 | VAQLVGYIAEHSAYHHGESSLQQTIINMAQTFVGSNNINFLEPCGQFGSR |
| 348 |  |  |
|  |  | \|\|\|\|\|\|\|\|\|\|\|\|\|\|\|\|\|\|\|\|\|\|\|\|\|\|\|\|\|\|\|\|\|\|\|\|\|\|\|\|\|\|\|\|\|\|\|\|\|\| |
| EMBOSS_001 | 301 | VAQLVGYIAEHSAYHHGESSLQQTIINMAQTFVGSNNINFLEPCGQFGSR |
| 350 |  |  |
| EMBOSS_001 | 349 | KEGGKDASAARYIFTKLASSTRSIFNEYDDPILKYLNEEGQKIEPQYYIP |
| 398 |  |  |
|  |  | \|\|\|\|\|\|\|\|\|\|\|\|\|\|\|\|\|\|\|\|\|\|\|\|\|\|\|\|\|\|\|\|\|\|\|\|\|\|\|\|\|\|\|\|\|\|\|\|\|\| |
| EMBOSS_001 | 351 | KEGGKDASAARYIFTKLASSTRSIFNEYDDPILKYLNEEGQKIEPQYYIP |
| 400 |  |  |
| EMBOSS_001 | 399 | VIPTILVNGCEGIGTGYSSFIPNYNYKDIIDNIKRYINKEPLIPMVPWYK |
| 448 |  |  |
|  |  | \|\|\|\|\|\|\|\|\|\|\|\|\|\|\|\|\|\|\|\|\|\|\|\|\|\|\|\|\|\|\|\|\|\|\|\|\|\|\|\|\|\|\|\|\|\|\|\|\|\| |
| EMBOSS_001 | 401 | VIPTILVNGCEGIGTGYSSFIPNYNYKDIIDNIKRYINKEPLIPMVPWYK |
| 450 |  |  |
| EMBOSS_001 | 449 | DFKGRIESNGKTGYETIGIINKIDNDTLEITELPIKKWTQDYKEFLEELL |
| 498 |  |  |
|  |  | \|\|\|\|\|\|\|\|\|\|\|\|\|\|\|\|\|\|\|\|\|\|\|\|\|\|\|\|\|\|\|\|\|\|\|\|\|\|\|\|\|\|\|\|\|\|\|\|\|\| |
| EMBOSS_001 | 451 | DFKGRIESNGKTGYETIGIINKIDNDTLEITELPIKKWTQDYKEFLEELL |
| 500 |  |  |
| EMBOSS_001 | 499 | TDEKHQLILDYIDNSSHEDICFTIKMDPAKLQKAEEEGLEKVFKLKSTLT |
| 548 |  |  |
|  |  | \|\|\|\|\|\|\|\|\|\|\|\|\|\|\|\|\|\|\|\|\|\|\|\|\|\|\|\|\|\|\|\|\|\|\|\|\|\|\|\|\|\|\|\|\|\|\|\|\|\| |
| EMBOSS_001 | 501 | TDEKHQLILDYIDNSSHEDICFTIKMDPAKLQKAEEEGLEKVFKLKSTLT |
| 550 |  |  |
| EMBOSS_001 | 549 | TTNMTLFDPNLKLQRYSTELDILKEFCYQRLKAYENRKSYLISKLEKEKR |
| 598 |  |  |
|  |  | \|\|\|\|\|\|\|\|\|\|\|\|\|\|\|\|\|\|\|\|\|\|\|\|\|\|\|\|\|\|\|\|\|\|\|\|\|\|\|\|\|\|\|\|\|\|\|\|\|\| |
| EMBOSS_001 | 551 | TTNMTLFDPNLKLQRYSTELDILKEFCYQRLKAYENRKSYLISKLEKEKR |
| 600 |  |  |
| EMBOSS_001 | 599 | IISNKTKFILAIVNNELIVNKKKKKVLVEELYRKGYDPYKDIN---KEEI |
| 645 |  |  |
|  |  | \|\|\|\|\|\|\|\|\|\|\|\|\|\|\|\|\|\|\|\|\|\|\|\|\|\|\|\|\|\|\|\|\|\|\|\|\|\|\|\|\|\|\| \|\|\|\| |
| EMBOSS_001 | 601 | IISNKTKFILAIVNNELIVNKKKKKVLVEELYRKGYDPYKDINKIKKEEI |
| 650 |  |  |
| EMBOSS_001 | 646 | FEQEL EDNEEIIAGITVKDYDYLLSMPIFSLTLEKVEDLLTQL |
| 688 |  |  |

| \|\|\|\|\| | | | \|\|\|\|\|\|\|\|\|\|\|\|\|\|\|\|\|\|\|\|\|\|\|\|\|\|\|\|\|\|\|\|\|\|\|\|\|\| |
| --- | --- | --- | --- |
| EMBOSS_001 700  EMBOSS_001 | 651  689 | FEQELLDAADNPEDNEEIIAGITVKDYDYLLSMPIFSLTLEKVEDLLTQL  KEKERELEILRNITVETMWLKDIEKVEEAIEFQRNVELSNREE------- | |
| 731 |  | \|\|\|\|\|\|\|\|\|\|\|\|\|\|\|\|\|\|\|\|\|\|\|\|\|\|\|\|\|\|\|\|\|\|\|\|\|\|\|\|\|\|\| | |
| EMBOSS_001 750 EMBOSS_001 | 701  732 | KEKERELEILRNITVETMWLKDIEKVEEAIEFQRNVELSNREESNHHHHH  - 731 | |
| EMBOSS_001 | 751 | H 751 | |

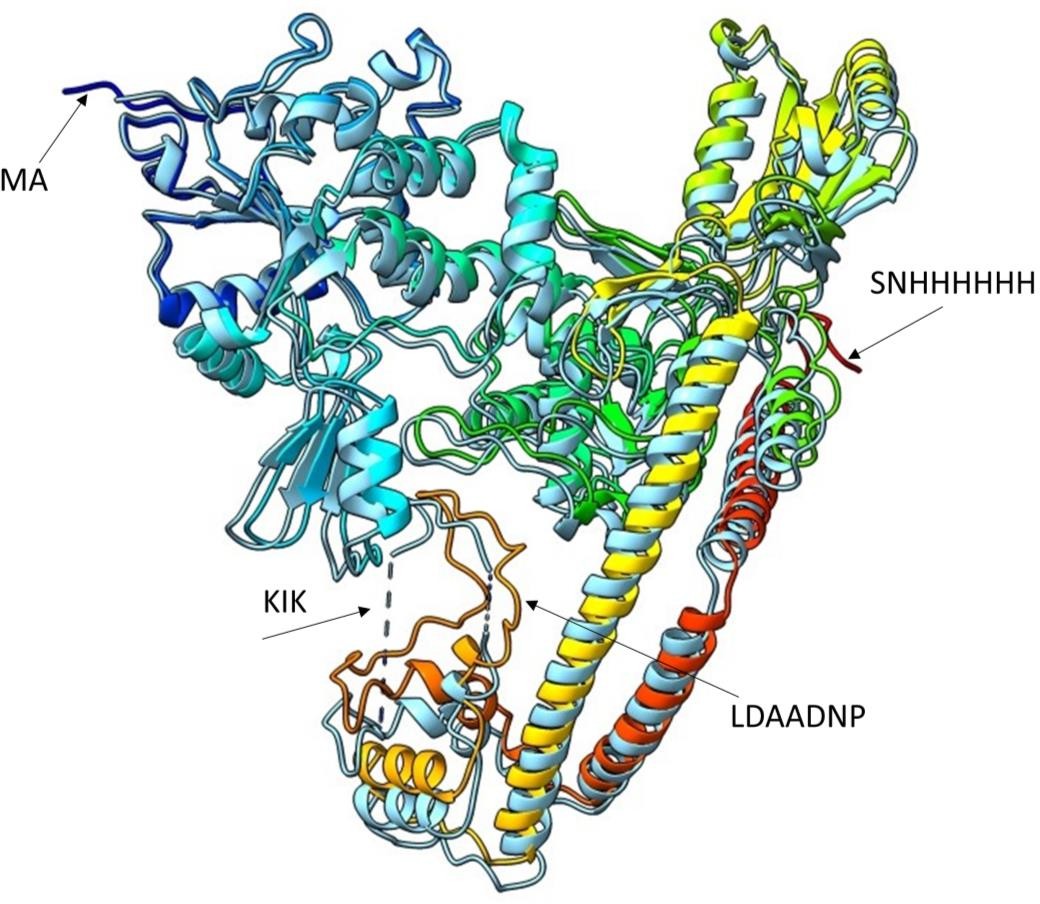

(a)

(b)

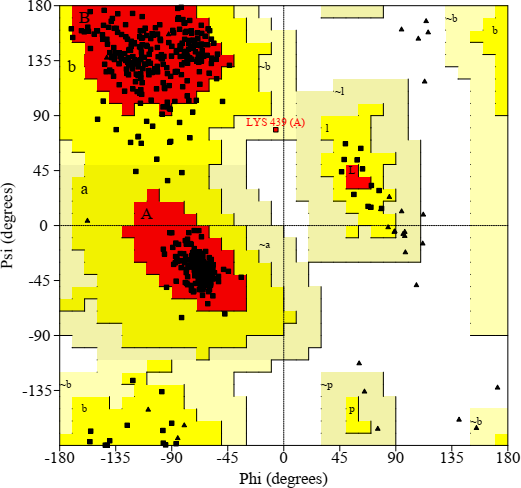

**Supplementary Figure S1**: (a) Modeled full length structure of Plasmodium falciparum DNA gyrase (6CA8) generated by using Modeller v10.6. The structure incorporates the missing amino acids through homology-based loop reconstruction. The backbone is demonstrated a ribbon diagram, color-coded from blue (N-terminal) to red (c-terminal). The replaced motifs are annotated: “MA” N-terminal motif, “KK” internal motif, “SNHHHHH” C-terminal polyhistidine, and “LDAADNP’ putative function motif, resulting in a complete and continuous protein model. The structure shows characteristic secondary elements, including alpha helices and beta sheets, as visualized in the final model. (b) Ramachandran plot of pfDNA gyrase model generated by PROCHECK, shows the distribution of backbone dihedral angles φ(phi) and ψ(psi) for all residues in the modeled protein structure.

| **S. No** | **Regions** | **Number of Residues** | **Percentage** |
| --- | --- | --- | --- |
| 1 | Residues in most Favored Regions (A, B, and L) | 635 | 92.6% |
| 2 | Residues in additional allowed regions (a, b, l, and p) | 50 | 7.3% |
| 3 | Residues in generously allowed regions (~a, ~b, and, ~p) | 1 | 0.1% |
| 4 | Residues in disallowed regions | 0 | 0.0% |
| 5 | Number of non-glycine and non-proline residues | **686** | **100%** |
| 6 | Number of end-residues (excl. Glycine and Proline) | 2 |  |
| 7 | Number of glycine residues (shown as triangles) | 42 |  |
| 8 | Number of proline residues | 21 |  |
|  | **Total number of residues** | **751** |  |

**Supplementary Table 1**: Ramachandran plot statistics of pfDNA gyrase. The table demonstrated the distribution of residues based on their backbone dihedral angles φ(phi) and ψ(psi) as analyzed by PROCHECK. Residues are shown in favored, additionally allowed, generously allowed, and disallowed regions according to steric constraints. The analysis includes non-glycine, non-proline residues, as well as glycine and proline residues separately due to their unique confirmational properties. The most favored regions indicate stereochemical reliable protein model with a high percentage of 92.6% residues.

| **S.No.** | **Compound name** | **Compound 2D structure** | **Smiles strings** | **Binding Affinity (kcal/mol)** | **Interacting residues** |
| --- | --- | --- | --- | --- | --- |
| **1** | Withanolide Q | 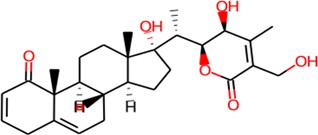 | OCC1=C(C)[C@@H]([C@@H](OC1=O)  [C@H]([C@@]1(O)CC[C@@H]2[C@] 1(C)CC[C@H]1[C@H]2CC=C2 [C@]1(C)C(=O)C=CC2)C)O | -8.57 | TYR385, ILE382, PRO381, TYR590 |
| **2** | Withanolide R | 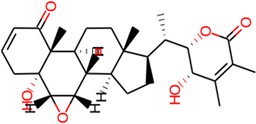 | O=C1O[C@H]([C@H](C(=C1C)C)O)[C@]  ([C@H]1CC[C@@H]2[C@]1(C)CC[C@H1 [C@H]2[C@@H]2O[C@@H]2[C@@]2 ([C@]1(C)C(=O)C=CC2)O)C | -8.61 | LYS392, LEU383, TYR385, LYS392, PRO381, ASP657, LYS567, GLU649 |
| **3** | Withanolide M | 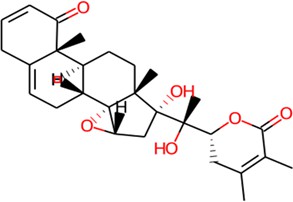 | CC1=C(C)C(=O)O[C@H](C1)[C@@]  ([C@@]1(O)C[C@H]2[C@@]3([C@] 1(C)CC[C@H]1[C@H]3CC=C3[C@]  1(C)C(=O)C=CC3)O2)(O)C | -8.69 | LYS392, GLU649, GLU648, LYS647, ILE645, ASN643 |
| **4** | Withanolide D | 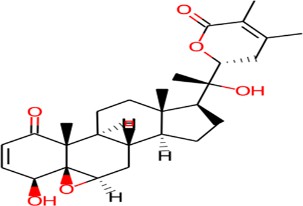 | CC1=C(C(=O)O[C@H](C1)[C@@](C)([C@ H]2CC[C@@H]3[C@@]2(CC[C@H]4[C@ H]3C[C@@H]5[C@]6([C@@]4(C(=O)C= C[C@@H]6O)C)O5)C)O)C | **-9.14** | ILE179, TRP177, LYS646, LEU656, LEU655, GLY176 |
| **5** | Withanolide E | 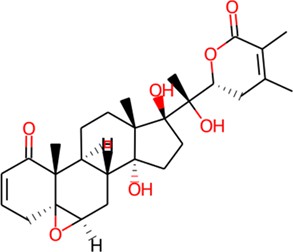 | CC1=C(C(=O)O[C@H](C1)[C@@](  C)([C@@]2(CC[C@@]3([C@@]2(C  C[C@H]4[C@H]3C[C@@H]5[C@]6(  [C@@]4(C(=O)C=CC6)C)O5)C)O)O  )O)C | **-9.73** | LYS392, LEU383, LYS647, GLU648, |
| **6** | Withanolide G | 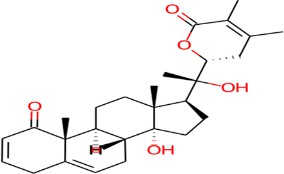 | CC1=C(C)C(=O)O[C@H](C1)[C@@]  ([C@H]1CC[C@@]2([C@]1(C)CC  [C@H]1[C@H]2CC=C2[C@]1(C)  C(=O)C=CC2)O)(O)C | -8.83 | LEU656, ALA658, THR172, ASN173 |
| **7** | Withanolide O 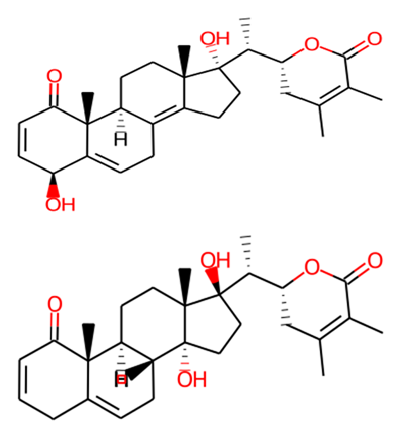 |  | CC1=C(C)C(=O)O[C@H](C1)[C@H]  ([C@@]1(O)CCC2=C3[C@H](CC  [C@]12C)[C@@]1(C)C(=O)C=C [C@@H](C1=CC3)O)C | **-9.00** | GLU652, ILE179, GLU389, GLN653 |
| **8** | Withanolide P |  | CC1=C(C)C(=O)O[C@H](C1)[C@H]  ([C@]1(O)CC[C@@]2([C@]1(C)CC[ C@H]1[C@H]2CC=C2[C@]1(C)C(=O) C=CC2)O)C | -7.89 | LEU165, ASN173, LYS646, ASP657 |

| **9** | Withanolide N | 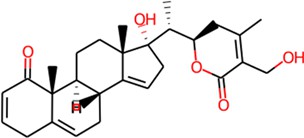 | OCC1=C(C)C[C@@H](OC1=O)[C@H] ([C@@]1(O)CC=C2[C@]1(C) CC[C@H]1[C@H]2CC=C2 [C@]1(C)C(=O)C=CC2)C | -8.16 | GLU654, LEU656, ASP657, ALA658 |
| --- | --- | --- | --- | --- | --- |
| **10** | Withanolide S | 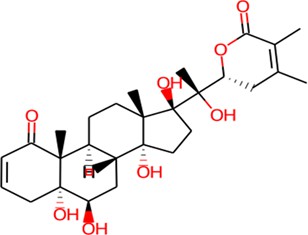 | CC1=C(C)C(=O)O[C@H](C1)[C@@]([C@]  1(O)CC[C@@]2([C@]1(C)CC[C@H]1[C@  H]2C[C@H]([C@@]2([C@]1(C)C(=O) C=CC2)O)O)O)(O)C | -8.04 | LYS647, PRO381, GLU648, ILE382, TYR385, LYS384 |
| **11** | Withanolide H | 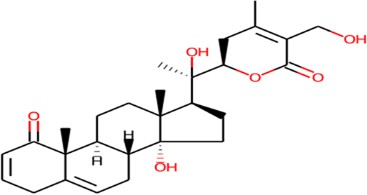 | OCC1=C(C)C[C@@H](OC1=O)[C@@] ([C@H]1CC[C@@]2([C@]1(C)CC[C@ H]1[C@H]2CC=C2[C@]1(C)C  (=O)C=CC2)O)(O)C | -7.45 | LEU383, TYR385, LYS392, ILE382 |
| **12** | Withanolide I | 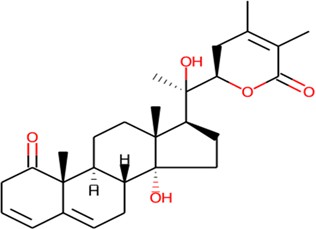 | CC1=C(C)C(=O)O[C@H](C1)[C@@]([  C@H]1CC[C@@]2([C@]1(C)CC[C@H]  1[C@H]2CC=C2[C@]1(C)C(=O) CC=C2)O)(O)C | -8.70 | LEU656, ASP657, ASN173, THR172 |

**Supplementary Table 2:** The table demonstrates the molecular docking results of various Withanolides derivatives along with SMILES strings revealed the steroidal lactone frameworks with variations in hydroxylation patterns and side chain modifications. The binding affinity(kcal/mol) indicates the predicted strength of interactions between withanolides and pfDNA gyrase (PDB:6CA8), with more negative values reflecting stronger binding energy followed by interacting residues.
